# Harnessing RNA-based DNA repair pathways for targeted gene editing

**DOI:** 10.1101/2024.04.09.588775

**Authors:** Nhan Huynh, Sin Kwon, Thomas A. McMurrough, Kurt W. Loedige, Marjan Tavassoli, Weijuan Shao, Heyuan Qin, Khanh Luu, Balpreet Dharni, Olha Haydaychuk, Brent E. Stead, David R. Edgell

**Affiliations:** Specific Biologics Inc., MaRS Discovery District, MaRS Centre, Toronto, ON, Canada; Department of Biochemistry, Schulich School of Medicine & Dentistry, Western University, London, ON, Canada

## Abstract

Recent studies have revealed a role for RNA in the repair of DNA double-strand breaks. Here, we show that the asymmetric DNA overhangs generated by the small TevSaCas9 dual nuclease informs a simple and robust editing strategy in human cells whereby Polθ and Rad52 are recruited to repair the double-strand break. The 2-nt, 3’ DNA overhang generated by the I-TevI nuclease domain of TevSaCas9 hybridizes with the 3’ end of a co-localized repair template guide RNA to specifically license repair. Substitutions that destabilize the repair duplex reduce editing efficiency. Targeted RNA-templated repair (rep-editing) harnesses cellular RNA-based DNA repair pathways to introduce precise nucleotide edits, deletions and insertions in human cells with high efficiency and fidelity independent of co-delivered repair functions. The small size of TevSaCas9 and RNA repair template offers delivery advantages over size-constrained or multi-component editing systems.

## INTRODUCTION

Repair of DNA damage is essential to cellular viability^1^. Site-specific programmable nucleases used for gene editing (editors) classically target specific sequences for modification by introducing DNA damage and that is subsequently repaired through recruitment of DNA-based repair pathways to install genomic edits^2–4^. However, a major challenge in the gene-editing field is the heterogeneous nature of repair events that impact the efficiency, fidelity, and spectrum of editing outcomes^5–7^. Recently developed strategies to enhance editing include Cas9-fusion proteins with added biochemical activities^8–10^, co-delivery of exogenous DNA polymerases or single-strand DNA repair templates^8, 11^, and knockdown of competing DNA repair pathways or over expression of repair factors^12–15^. These strategies come at the cost of increasing the number, size, complexity and potential toxicity of components that need to be delivered to cells.

In mammalian systems, double-strand breaks (DSBs) are repaired through well-characterized pathways that are grouped into homology dependent repair (HDR) and non-homologous mediated end-joining (NHEJ) pathways^17^. A role for RNA in the repair of cellular DSBs is gaining appreciation through studies in yeast and human cell lines showing that homologous RNA transcripts can mediate DSB repair in expressed genes^18,19^and through direct repair of DSBs by RNA oligonucleotides^20^. Rad52 is required for RNA-mediated DNA repair by facilitating pairing between homologous RNA and DNA through an inverse strand exchange activity^18,21^. The translesion DNA polymerase *θ* (Pol*θ*) that participates in alternative-end joining repair (also called microhomology-mediated end-joining or theta-mediated end joining)^22, 23^, exhibits high fidelity as a reverse transcriptase (RT) on RNA-DNA templates and can stimulate repair of model DSB substrates with an RNA repair template in HEK293 cells^24^; similar RNA-repair functions are found for PolV and Pol*η*^25, 26^. These observations suggest that RNA could act as an efficient template for the repair of targeted DSBs through recruitment of endogenous repair functions if an RNA repair template could be localized to the break site.

We previously described TevCas9^7, 27^, a small, chimeric RNA-guided editor that generates two DSBs with incompatible DNA ends, a Tev 2-nt 3’ overhang^28^ and a Cas9 blunt end. TevCas9 is a fusion of the GIY-YIG nuclease domain and interdomain linker from the homing endonuclease I-TevI (hereafter, Tev)^29^ to Cas9 from Streptococcus pyogenes (SpCas9) or Staphylococcus aureus (SaCas9) with a coding size of <3.5-kb in the case of the TevSaCas9 fusion (27) (Fig. 1A) The Tev nuclease domain sequentially nicks the bottom (*↑*) and top strands (*↓*) at preferred 5’-CN*↑* NN*↓* G-3’ motifs^30, 31^ that are positioned upstream of the 5’ end of the Cas9 gRNA binding site.

**Figure 1.**
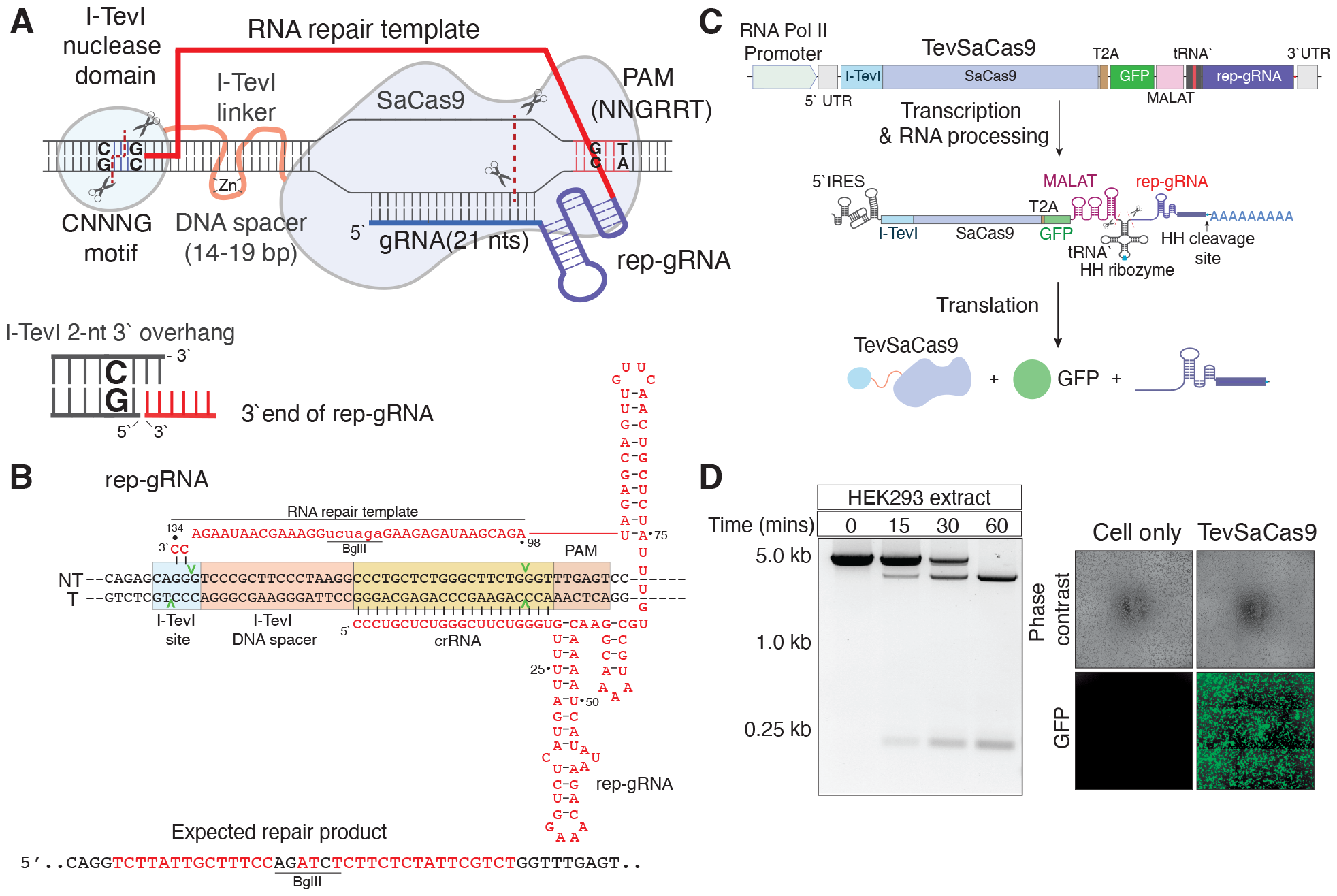
Rep-editing design and expression. **A**, Schematic of rep-editing outlining a TevSaCas9 target site with the individual DNA targeting components, the domain structure, and the rep-gRNA. The I-TevI linker zinc finger is indicated by yellow dot. Cleavage by the I-TevI domain leaves a 2-nt, 3’ overhang that is complementary to the 3’ end of the rep-gRNA, while cleavage by SaCas9 generates a blunt DNA end. **B**, The AAVS1 target site, expected repair product, and structure and interactions of the rep-gRNA with the target site. **C**, Schematic and predicted processing of the all-in-one construct with individual components indicated (not to scale). RNA Pol II promoter, mini CMV promotoer; MALAT, RNA stability element from the MALAT1 non-coding RNA; HH, hammerhead ribozyme; T2A, self-splicing peptide linker; IRES, internal ribosome entry site^16^. **D**, Left, *in vitro* processing of the all-in-one mRNA transcript by HEK293 cell extracts, incubated for the indicated time and resolved on an 1.5% agarose gel. Right, eGFP activity in HEK293 cells transfected with the all-in-one construct.

Here, we show that the directional nature of the TevSaCas9 cleavage products informs a simple and highly efficient editing strategy using a co-localized modified repair gRNA template (rep-gRNA) (Fig. 1A and 1B) that hybridizes to the Tev 2-nt 3’ overhang to prime repair through the endogenous RNA repair activities of Pol*θ* and Rad52. TevSaCas9 RNA-templated editing (rep-editing) can install nucleotide changes, deletions and insertions with high efficiency and fidelity. The small sizes of TevSaCas9 and rep-gRNA are easily accommodated by a single AAV viral delivery vector, offering advantages over size-constrained and multiple-component editing modalities, and representing a different approach to gene editing that uses RNA-templated repair of localized DSBs.

## RESULTS

### A modified gRNA for targeted repair in human cells

To test directional, RNA-templated repair using the TevSaCas9 nuclease, we created a modified guide RNA by fusing an RNA repair sequence to the 3’ end of standard SaCas9 single guide RNA (sgRNA)^32^ composed of the gRNA portion that is the complement of the target site and the tracrRNA portion to create a rep-gRNA (repair templated gRNA) (Fig. 1A and 1B). We envisioned a repair scenario where the 3’ end of the rep-gRNA was the complement of the 2-nt 3’ overhang generated by the Tev nuclease domain. These complementary bases would form a duplex RNA-DNA hybrid with a 2-nt 3’ overhang and act as a priming site for repair (Fig. 1A), possibly through reverse transcription of the RNA repair template. The location of the targeted edit can be at any position between the Tev and SaCas9 cleavage sites up to the rep-gRNA scaffold region. We first designed a rep-gRNA where the gRNA portion targeted a site in the *AAVS1* safe harbour locus^33, 34^ (Fig. 1B, Table S1). The *AAVS1* rep-gRNA contained a repair sequence of 37 nucleotides in length that was dissimilar to the *AAVS1* sequence and included a diagnostic BglII site. Crucially, the 3’ end of the rep-gRNA possessed two complementary nucleotides (5’-CC-3’) to the 2-nt 3’ overhang left by Tev cleavage at the appropriately positioned 5’-CA*↑*GG*↓*G-3’ site (the 2-nt 3’ overhang product is 5’-GG-3’) (Fig. 1B).

To confirm activity of TevSaCas9 and SaCas9 (Fig. S1A and S1B) at the *AAVS1* site, we used a standard single guide gRNA (sgRNA) targeted to this site and demonstrated robust activity *in vitro* and *in vivo* activity in HEK293 cells for TevSaCas9 (Fig. S1C and S1D). We also determined that the optimal spacing of the CNNNG Tev cleavage motif was 20- or 30-bp upstream of the SaCas9 binding site, consistent with spacing preferences for the native Tev nuclease and other Tev-based chimeric nucleases^30, 35–37^ (Fig. S2A and S2B). We estimate that *∼*75% of SaCas9 sites in the human genome, defined by the presence of an NNGRRT PAM site and 21-nt gRNA binding site, possessed appropriately spaced CNNNG motifs that could support Tev cleavage (Fig. S2C).

To deliver TevSaCas9 and the rep-gRNA to cells, we created an all-in-one format construct where a single RNA polymerase II transcript of ∼ 4.5 kb contains the TevSaCas9 coding sequence and rep-gRNA and that would easily be accommodated by the coding capacity of a single AAV viral vector (Fig. 1C). This setup would avoid the requirement for a separate RNA polymerase III promoter to express the gRNA. Since this setup removed a poly(A) tail from the TevSaCas9 coding sequence, we included a MALAT1 sequence at the 3’ end of TevSaCas9 for stability^38^. To stimulate post-transcriptional processing of the all-in-one mRNA, we inserted an alanine tRNA sequence (ala tRNA) between the TevSaCas9-eGFP coding region and rep-gRNA, as well as a *trans*-acting hammerhead (HH) ribozyme sequence in the hairpin of the alanine tRNA to act on a HH cleavage site located between the 3end of the rep-gRNA and the poly(A) signal. To confirm correct processing, we incubated *in vitro* transcribed TevSaCas9-eGFP/rep-gRNA all-in-one mRNA with a HEK293 cell extract, and observed processing into the two predicted products (TevSaCas9-eGFP and rep-gRNA) (Fig. 1D). We also transfected HEK293 cells with the TevSaCas9-eGFP/rep-gRNA all-in-one mRNA and observed robust eGFP activity (Fig. 1D).

### Rep-editing in human cell lines

The TevSaCas9/rep-gRNA all-in-one construct targeting the *AAVS1* safe harbour site was introduced into HEK293 cells by four different methods; AAV2 transduction, lipid nanoparticles containing mRNA, plasmid DNA (pDNA) transfection, or a ribonucleoprotein complex (RNP) consisting of purified TevSaCas9 complexed with *in vitro* transcribed rep-gRNA (Fig. 2A). We did not select or enrich for transfected cells and thus the reported editing rates are determined from DNA isolated from total cells in any given experiment. We observed robust editing at the AAVS1 site with all four delivery modalities by BglII digests (56 +/- 12%) and by deep sequencing (52 +/- 14%) of target sites PCR amplified from genomic DNA (Fig. 2B). AAV2 delivery resulted in the highest editing rates (67 +/- 10%) whereas the editing rate for RNP transfections was lowest (43 +/- 11%) (Fig. 2C), likely reflecting lower RNP transfection versus AAV2 transduction efficiency.

**Figure 2.**
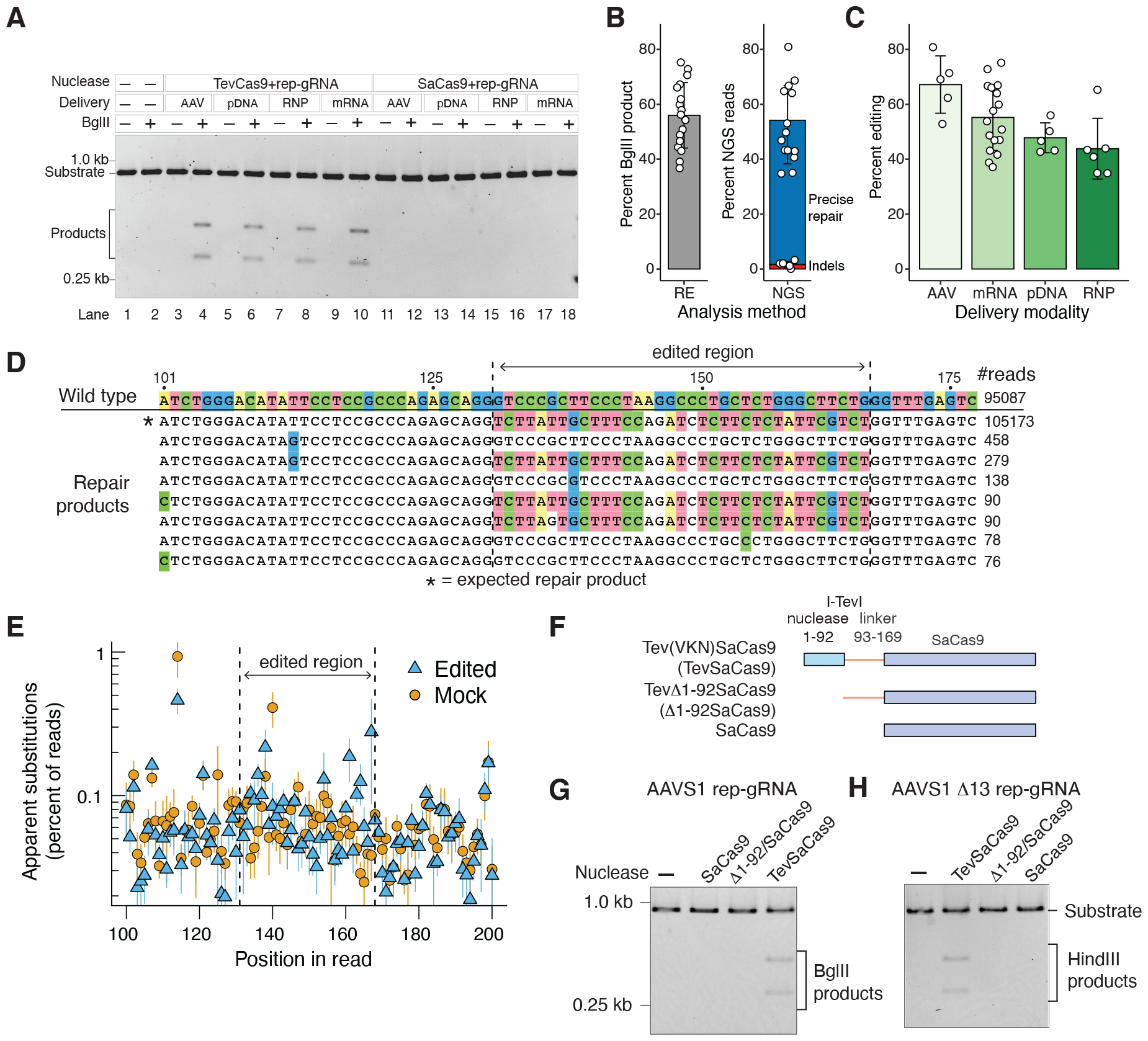
Rep-editing at the *AAVS1* safe harbour site. **A**, BglII digestions of *AAVS1* genomic target sites PCR amplified from treated HEK293 cells. AAV; adeno-associated virus 2; pDNA, plasmid DNA; RNP, ribonucleoprotein particle of TevSaCas9 and rep-gRNA; mRNA, all-in-one construct. **B**, Editing outcomes in total cells at the *AAVS1* site by analysis method; RE, BglII restriction digest; NGS, next generation sequencing. **C**, Editing outcomes by delivery method. Barplots are the mean of all replicates with whiskers representing the standard deviation from the mean. Dots are individual replicates. **D**, Example reads from deep sequencing of the *AAVS1* site in TevSaCas9/rep-gRNA treated cells. The number of reads for each sequence is indicated on the right. Differences relative to the wild-type sequence are colored by nucleotide. **E**, Plot of apparent nucleotide substitutions from deep sequencing of *AAVS1* PCR amplicons from edited cells (blue triangles, 6 replicates) or mock transfected cells (orange circles, 4 replicates). Points are mean values with whiskers representing the standard deviation from the mean. **F**, Schematic of TevSaCas9 constructs with the Tev nuclease domain and linker regions indicated (not to scale). VKN, V117F, K135R, N140S substitutions in the Tev linker domain that increase activity on sub-optimal DNA spacer sequences^7^. **G**, Editing in HEK293 cells transfected with the *AAVS1* rep-gRNA and the indicated constructs. Shown is a BglII digest of PCR amplified *AAVS1* target site amplified from total cells, with BglII products indicated. **H**, As in panel **G** but with the *AAVS1* rep-gRNA designed to insert a 13-bp deletion. Successful editing creates a HindIII site.

Using PCR primers that would distinguish correctly from incorrectly oriented editing events, we showed that TevSaCas9/rep-gRNA editing generated correctly orientated repair products (Fig. S3A). Analysis of deep sequencing reads from 6 independent TevSaCas9-rep-gRNA experiments confirmed both the directional insertion and fidelity of the editing, with 52 +/- 14% of reads corresponding to the expected repair product with no other templated nucleotide changes (Fig. 2D and 1E). Less than 0.1% of reads corresponding to incorrect repair products, while 1.6 +/- 1.1% of reads possessed indels. No reads were identified where repair extended beyond the RNA repair template region to include sequence in the scaffold region of the rep-gRNA, indicating that editing and repair is specified by only the RNA repair template portion of the rep-gRNA. Moreover, apparent nucleotide substitutions in sequencing reads for edited cells were indistinguishable from those in mock treated cells (Fig. 2E). We also designed a rep-gRNA to introduce a 13-bp deletion in the *AAVS1* site (*AAVS1* Δ13) that would create a HindIII site (Fig. S3A, Table S1). We found 44% of deep sequencing reads were consistent with a precise 13-bp deletion and 4.1% of reads were consistent with other deletion lengths, as compared to 1.4% of sequencing reads from mock treated cells possessing length differences (Fig. S3B and S3C).

To extend our results to different target sites and cell types, we designed rep-gRNAs to correct clinically relevant mutations in the *CFTR* gene in 16HBE cells that are individually engineered to contain the ΔF508, G542X and W1282X mutations (Fig. S5A-C, Table S1, Supplementary Text)^39^. We also designed a rep-gRNA to correct the *SERPINA1* E342K mutation that is causative for α1 antitrypsin (A1ATD) deficiency using an A1ATD primary fibroblast cell line^40^ (Fig. S5D). These mutations are representative of three common types of therapeutic edits; insertions (ΔF508), transversions (G542X and E342K) and transitions (W1282X). In addition, the *CFTR* G542X, W1282X and *SERPINA1* E342K sites contain bystander bases that can be problematic for base editing^41^. Using four different analysis methods on DNA isolated from total cells transfected with TevSaCas9/rep-gRNA mRNAs, we found robust editing at all sites (31 +/- 7% for ΔF508, 51 +/- 8% for G542X, 49 +/- 7% for W1282X, and 42 +/-13% for E324K) (Fig. S5E and S5F). Deep sequencing of editing at the *CFTR* G542X and *SERPINA1* E342K sites revealed precise repair with no evidence of non-templated (or bystander) nucleotide changes.

### Rep-editing requires the Tev nuclease domain

To specifically test the requirement for the Tev nuclease domain in rep-editing, we deleted residues 1-92 (Δ1-92) that correspond to the structurally defined nuclease domain while retaining linker residues 93-169 (Fig. 2F). No repair at the *AAVS1* site by BglII digestion, or at the *AAVS1* Δ13 site by HindIII digestion, was observed when the Δ1-92 all-in-one mRNA construct was lipofected into HEK293 cells (Fig. 2G and 2H) as compared to the TevSaCas9 WT construct (Fig. 2A lanes 2-10 and Fig. S4B). We repeated these experiments with wild-type SaCas9 in place of TevSaCas9 (Fig. 2A, lanes 11-18; Fig. 2G and 2H). No repair was observed with SaCas9/rep-gRNA at the *AAVS1* site. Purified SaCas9 (Fig. S1a) complexed with *in vitro* transcribed rep-gRNA was active *in vitro* on a PCR generated *AAVS1* target (Fig. S1E), implying that the lack of observed repair in HEK293 cells is not due to SaCas9 inactivity.

Collectively, this data shows that targeted DSBs in human cells can be repaired by an RNA-templated process (rep-editing). Repair is efficient and has high fidelity at multiple sites in three different cell types. Moreover, rep-editing is dependent on a Tev-induced DSB upstream of the SaCas9 gRNA binding site.

### Increasing rep-gRNA 3’ identity enhances editing

To further probe the impact of nucleotide identity between the 3’ end of the rep-gRNA and Tev cleavage site on repair efficiency, we designed a series of *AAVS1* targeting rep-gRNAs where the length of complementarity to sequence upstream of the Tev cleavage site was increased in 2-nt increments to a maximum of 18 nts (Fig. 3A; we termed these 3’ overlaps). We also varied complementarity towards the PAM site from the SaCas9 cleavage site to a maximum of 6 nts (Fig. 3A; we termed these 5’ overlaps). To enable analysis of multiple designs, we used a lipofection strategy with an mRNA encoding TevSaCas9 and *in vitro* transcribed rep-gRNAs and used quantitative reverse-transcription PCR to measure the ratio of correctly repaired sequence compared to unrepaired sequence in *AAVS1* transcripts in each sample^42^. We found that 2-nt of complementarity to the Tev 2-nt overhang at the 3’ end of the rep-gRNA was essential for repair and that complementarity beyond 18-nts reduced the efficiency of repair (Fig. 3B, dark green bars). We found similar increases in efficiency with deep sequencing (Fig. S6A). The fidelity of editing was not impacted for the 3’ overlapping rep-gRNAs (Fig. S6B). In contrast, there was no increase in repair with 5’ overlap rep-gRNAs that had increasing complementarity PAM proximal to the Cas9 cut site (Fig. 3B). We also considered the possibility that the 3’ end of the rep-gRNA would be susceptible to degradation by 3’ exonucleases or that auto-inhibitory structures in the rep-gRNA could impact repair efficiency, as has been suggested with prime editing pegRNAs^43^. To test this, we included a protecting oligonucleotide complementary to the length of the repair template. When we repeated the lipofections with the protected rep-gRNAs, we observed we observed a modest 1.4 1.6-fold increase in repair efficiency for rep-gRNAs with 12, 14 or 16 nt 3’ overlaps (Fig. 3C, light green bars).

**Figure 3.**
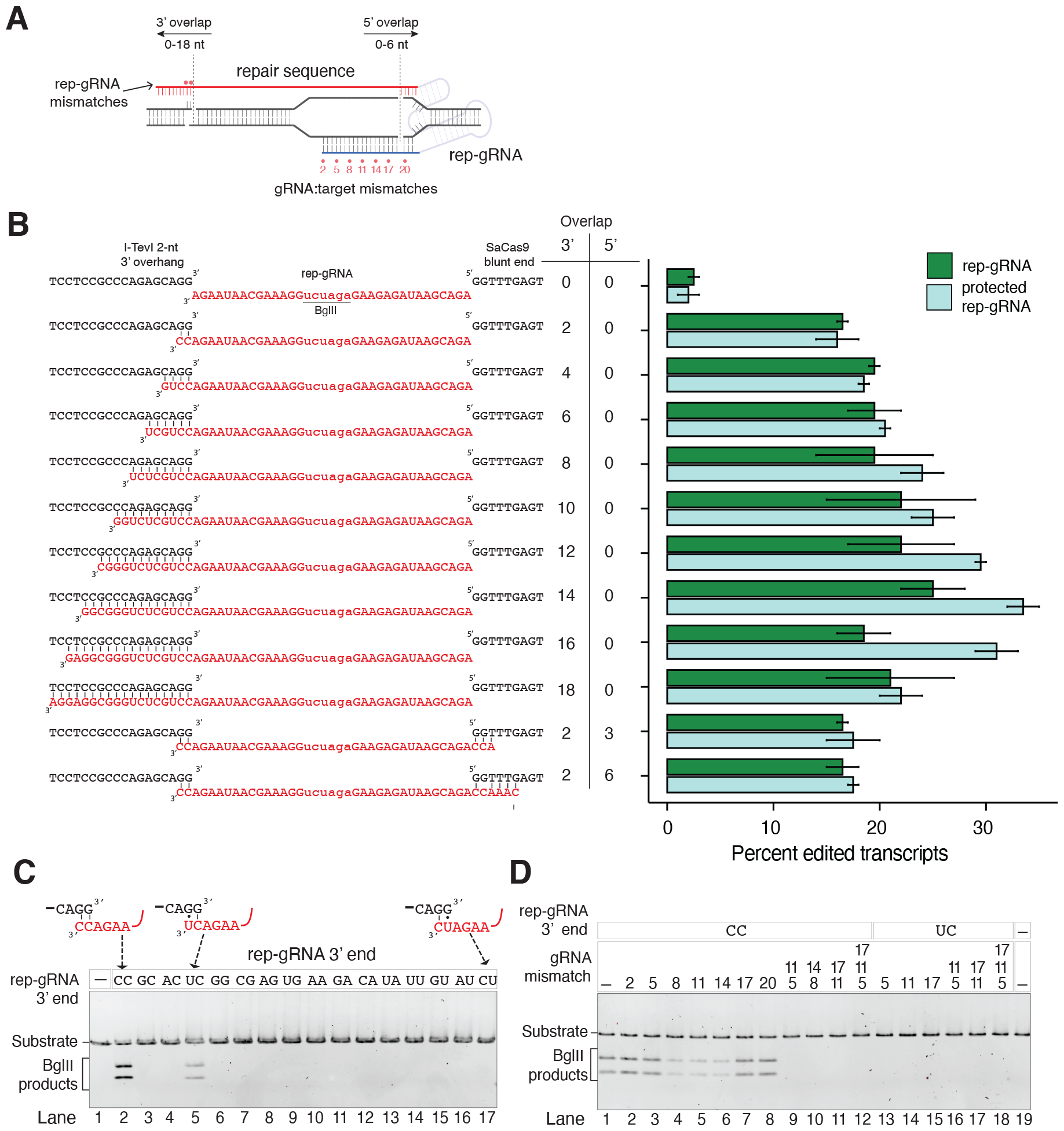
A stable RNA:DNA hybrid is required for rep-editing. **A**, Schematic of rep-gRNAs used to test base pairing between the 3’ end of rep-gRNA and Tev overhang at the AAVS1 site. **B**, Impact on editing efficiency of rep-gRNAs with different length 3’ and 5’ overlaps. The barplots are the ratio of repaired to unrepaired sites as judged by quantitative-PCR of the AAVS1 target site in treated cells. Dark green bars are cells treated with TevSaCas9/rep-gRNA while light green bars are cells treated with protected rep-gRNA. **C**, Impact of mismatches at the terminal two nucleotides of the rep-gRNA on editing. Shown is a representative gel of BglII digests of AAVS1 target site amplicons from treated HEK293 cells. **D**, Impact of mismatches between the gRNA portion of the rep-gRNA on repair at the AAVS1 site with a rep-gRNA with an exact match to the Tev overhang (CC) or a wobble base pair (UC). Nucleotide mismatches in the gRNA relative to the target site are indicated by position above the gel image.

We also probed the tolerance to mismatches between the 3’ end of the rep-gRNA and the Tev overhang, as well as mismatches to the *AAVS1* target site (Fig. 3A, 3C, and 3D). We found that apart from the exact 2-nt match between the 3’ rep-gRNA (5’-CC-3’) and Tev overhang (5’-GG-3’) (Fig. 3C, lane 2), the only other rep-gRNA that promoted repair had a 5’-CU-3’ sequence that would create an A-U wobble at the 3’ end (Fig. 3C, lane 5). Interestingly, the rep-gRNA with a 5’-UC-3’ sequence that would create at G-U wobble one base internal to the 3’ end did not support repair (Fig. 3C, lane 17). Single nucleotide mismatches to the *AAVS1* site in the rep-gRNA also supported repair but only with a rep-gRNA with a 5’-CC-3’ sequence (Fig. 3D, lanes 1-8) and not with a 5’-UC-3’ sequence (Fig. 3D, lanes 13-19). Double or triple nucleotide mismatches to the *AAVS1* site did not support repair with either rep-gRNA (Fig. 3D, lanes 9-19). We extended this analysis by profiling off-target cleavage at 38 computationally predicted sites for the *AAVS1* gRNA that had 4 or fewer mismatches to the gRNA as well as CNNNG motifs that were spaced at various positions upstream of the gRNA binding site (Supplemental Text, Fig. S7 and S8). We did not observe significant levels of editing at the off-target sites, whereas robust on-target editing (between 40-80%) was found.

Collectively, this data shows that stability between the 3’ end of the rep-gRNA and Tev cleavage overhang directs repair and that hybridization of a protecting oligonucleotide to the rep-gRNA can modestly increase repair. Our results suggest that the TevSaCas9/rep-gRNA has at least three distinct layers of specificity; identity between the gRNA portion of the rep-gRNA and target site; complementarity between the 3’ end of the rep-gRNA and the CNNNG motif (CAGGG in the case of the AAVS1 site); and spacing of the Tev cleavage motif from the gRNA binding site.

### Pol*θ* and Rad52 facilitate rep-editing

To better define the cellular repair mechanisms involved in rep-editing, we transduced HEK293 cells with AAV2 encoding TevSaCas9 and the *AAVS1* rep-gRNA in the presence of increasing concentrations of the small molecule inhibitors SCR7, an inhibitor of DNA ligase IV^44^; B02, an inhibitor of Rad51^45^; D-103, an inhibitor of Rad52^46^; and ART558, an inhibitor of Pol*θ* ^47^ (Fig. 4A). After 24-hr growth post transduction, the *AAVS1* target site was amplified and digested with BglII to assess the extent of repair. We observed diminished rep-editing with the D-103 and ART558 inhibitors (Fig. 4A). To test the specificity of each inhibitor to knockdown the relevant DNA repair pathway, cells treated with each inhibitor also received AAV2 encoding TevSaCas9 and a standard gRNA targeting the *AAVS1* site co-delivered with a lipofected duplex DNA with 40-bp homology arms (HDR dsDNA), a dsDNA donor with no homology arms (NHEJ dsDNA), or single-stranded RNA repair template (ssRNA) (Fig. 4A). As anticipated, inhibition of repair was specific for each inhibitor; SCR7 inhibited NHEJ repair with a dsDNA template; B02 inhibited HDR with the dsDNA template with homology arms; and D-103 inhibited ssRNA repair. To confirm the involvement of Pol*θ* and Rad52 in rep-editing, we transfected HEK293 cells with mRNA encoding TevSaCas9 or SaCas9, each with a C-terminal hemagglutinin tag (HA) for immunoprecipitation, and the rep-gRNA targeting *AAVS1*. Both TevSaCas9 and SaCas9 were present in extracts after immunoprecipitation with anti-HA magnetic beads (Fig. 4B and Fig. S5). We only detected the presence of Pol*θ* in extracts from TevSaCas9 transfections with an anti-Pol*θ* antibody, whereas Rad52 was detected with an anti-Rad52 antibody in extracts from both TevSaCas9 and Cas9 transfected cells. Taken together, these data suggest that RNA templated rep-editing is specific for TevSaCas9/rep-gRNAs and mediated by Rad52 and Pol*θ* through an RNA-templated repair pathway and not through NHEJ or HDR DNA repair pathways (Fig. 4C).

**Figure 4.**
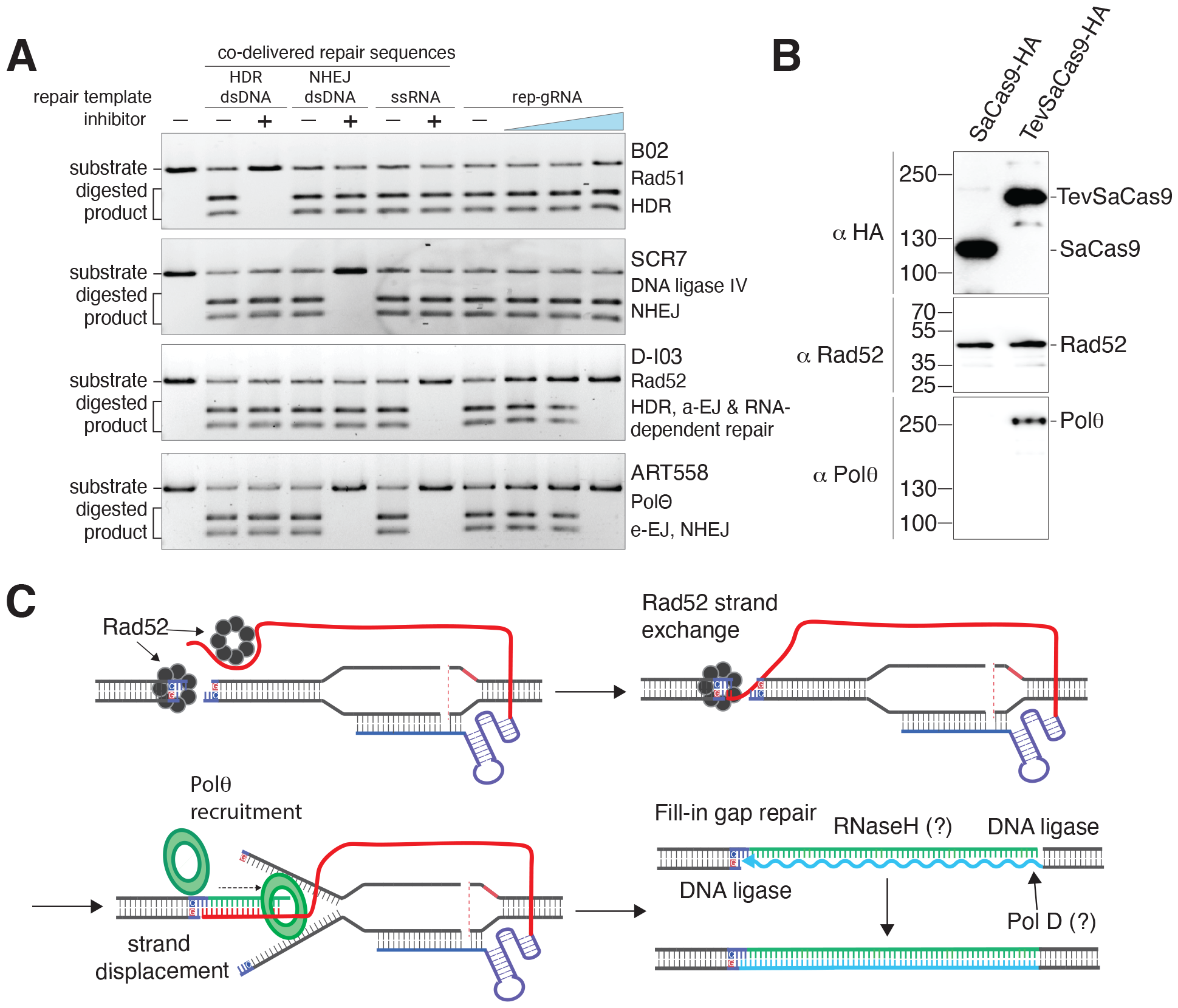
Pol*θ* and Rad52 mediate rep-editing. **A**, Small molecule inhibition of repair. On the right, HEK293 cells were treated with the B02, SCR7, D-103 or ART558 inhibitors at the same time as transduction with an AAV2 encapsidating TevSaCas9 and the rep-gRNA. Increasing concentration of inhibitor is indicated by a blue triangle. *AAVS1* target site was PCR amplified and digested with BglII. On the left, editing as determined by BglII digestion of amplified *AAVS1* target sites in cells transduced with AAV2-TevSaCas9 and different co-lipofected repair sequences as indicated, along with the indicated small molecule inhibitors. **B**, Co-immunoprecipitation with an anti-HA antibody of extracts from HEK293 cells transfected with mRNA TevSaCas9/rep-gRNA or SaCas9/rep-gRNA, followed by western blot with an anti-Pol*θ* or anti-Rad52 antibody. Sizes (in kd) are indicated on the left of the gel image. **C**, Model for rep-editing. Cleavage by TevSaCas9/rep-gRNA recruits Rad52 that can bind to RNA, or DNA, or mediate RNA:DNA strand exchange at the Tev cleavage site. Formation of a hybrid RNA:DNA structure with a 3’-OH recruits Pol*θ*. Resolution of the repair intermediate by a fill-in gap repair process involving DNA ligase and an unidentified DNA polymerase.

## DISCUSSION

Here, we show that rep-editing is a simple, robust and versatile method of gene editing. The high efficiency of rep-editing derives from the co-localization of the RNA repair template and DSB by genetically encoding the repair template and gRNA as a single unit. This strategy offers advantages in terms of simplicity and delivery over other methodologies that co-localize a DNA repair template and Cas9^8, 48, 49^. Rep-editing is also differentiated from size-constrained or multi-component Cas9-editing strategies in a number of ways. These include: its small size that is easily accommodated by a single AAV vector and other delivery modalities; recruitment of endogenous repair functions that negates co-delivery of accessory factors and repair templates; and the ability to introduce insertions, deletions or nucleotide changes through a flexible rep-gRNA design. More generally, our data also provides further support of a role for RNA in the repair of DNA damage in human cells.

Rep-editing uses a co-localized RNA repair guide to stimulate repair through an RNA-templated pathway that recruits the endogenous RNA-repair functions of Pol*θ* and Rad52. The DNA:RNA hybrid formed by pairing of the Tev 3’ overhang with the 3’ end of the rep-gRNA mimics the lengths of DNA microhomologies that occur at DSBs and that function to recruit Pol*θ* ^22, 23, 50–52^. The observed fidelity of rep-editing is consistent with the low substitution rates of Pol*θ* on RNA-DNA templates^24^. RNA-templated DNA repair was shown to involve the Rad52 RNA-DNA strand exchange activity^21^ which we envision mediates pairing of the 3’ Tev overhang with the 3’ end of the rep-gRNA, particularly for rep-gRNAs that have longer 3’ overlaps that extend beyond the 2’-nt Tev overhang.

RNA-templated repair in rep-editing is similar to the target-primed reverse transcription (TPRT) mobility pathway of non-LTR retrotransposons and group II introns where a DNA endonuclease-RT fusion nicks the DNA target site to initiate repair using the RNA of the mobile element as a template^53, 54^. The recently developed PRINT technology uses the TPRT mechanism to deliver transgenes of up to 4kb to safe harbour sites in human cells^55^. Prime editing with a Cas9 nickase-RT fusion also parallels TPRT as the Cas9-generated DNA nick and a modified gRNA template (pegRNA) prime reverse transcription to install genomic edits^9^. Interestingly, prime editing efficiency can be enhanced by using a second gRNA to introduce a nick in the non-edited strand^9^, essentially creating a staggered DSB to stimulate repair. There are important design differences between rep-gRNAs described here and pegRNAs. Notably, the repair template portion of rep-gRNAs can be longer than the corresponding primer-binding site and repair template portion of pegRNAs because the 3’ end of the rep-gRNA that hybridizes with the upstream Tev overhang is not complementary to the gRNA portion of the rep-gRNA, as is the case with pegRNAs. This design feature of rep-gRNAs presumably reduces auto-inhibitory structures that could impact editing efficiency. At this point, we do not know the upper limit of insertions that are possible with rep-editing, although a single AAV vector can deliver a genetically encoded TevSaCas9 and rep-gRNA with up to a 1-kb repair template, while lipofection with co-packaged TevSaCas9 mRNA and *in vitro* transcribed rep-gRNA could allow for longer repair templates.

Classical approaches to gene editing were largely based on repair of DSBs. Current Cas9 gene editors use different strategies to install genomic edits due to concerns that Cas9 off-target DSBs will be repaired by mutagenic pathways to introduce unwanted edits. In this regard, two aspects of rep-editing mitigate off-targets and contribute to the high fidelity and specificity of on-target editing. First, the sequence requirements of the TevSaCas9 nuclease create a target site of ∼ 38-bp. The optimal spacing of 15-25 bp between the Tev cleavage site and the gRNA binding site further contributes to specificity because not all off-target gRNA sites are permissive for Tev cleavage due to sub-optimal spacing. Our data show that while the SaCas9/rep-gRNA complex can cleave DNA substrates *in vitro*, we see no evidence of repair in cells transfected with SaCas9/rep-gRNA likely because the rep-gRNA is not designed to to hybridize with sequence surrounding the SaCas9 cleavage site to initiate *in vivo* repair. Second, the requirement for base pairing between the 3’ end of the rep-gRNA and 2-nt 3’ Tev overhang licenses specific repair by providing a priming site for Pol*θ*. For instance, the *AAVS1* target site possesses an alternative 5’-CAGAG-3’ site immediately upstream of the on-target 5’-CAGGG-3’ site. Similar alternative 5’-CNNNG-3’ motifs are also found at the *CFTR* G542X and *SERPINA1* E342K sites, and at the *AAVS1* rep-gRNA off-target sites. However, we found no evidence of cleavage and repair at the alternative Tev sites in deep sequencing of edited cells because in all cases the 3’ end of the rep-gRNA was designed to be a complement of the on-target Tev motif. If Tev is cleaving at alternative 5’-CNNNG-3’ sites in cells, the rate of cleavage and repair must be below the detection limit our deep sequencing experiments, or the cleaved sites are substrates for faithful NHEJ repair.

A number of questions related to rep-editing remain unanswered yet could stimulate future improvements. We do not know if rep-editing can utilize DNA nicks instead of a DSB made by the Tev domain to stimulate repair. The Tev nuclease and related isoschizomers^56–58^ function as a monomers to use a single active site to sequentially nick DNA on each strand to generate a DSB. Attempts to isolate Tev nicking variants have been unsuccessful. The Tev domain requires a CNNNG cleavage motif but not all NNN triplets are equally preferred^36, 59^. Previously isolated up-activity mutants in the Tev nuclease and linker domains increase cleavage on some NNN triplets^7, 59^ and could enhance editing at sub-optimal sites. Alternatively, GIY-YIG orthologs with different cleavage requirements^60, 61^ could be substituted into the TevSaCas9 architecture to broaden the targeting range in rep-editing. It is also possible that unrelated nuclease domains with different cleavage preferences and products (5’ versus a 3’ overhang) could be substituted for Tev, although the domains would need to function as monomers given the current protein architecture. Determining rep-editing activity and specificity in non-dividing cells would enable a wide-range of applications. Genome-wide off-target profiling *in vivo* would provide further insight into the specificity of rep-editing, and would be a prerequisite for the editing of therapeutically relevant target sites by RNA-templated repair of targeted DSBs.

## Acknowledgements

NSERC-MITACS Alliance grant ALLRP/565307-2021 (DRE, BES); Cystic Fibrosis Foundation Therapeutic Development Award SPECIFIC21W0-SC (Specific Biologics Inc.)

## Competing interests

B.E.S. is founder and employee of Specific Biologics Inc. D.R.E is co-founder of Specific Biologics Inc and is a member of the Scientific Advisory Board. N.H., T.A.M, S.K., and M.T. are employees of Specific Biologics Inc.

## Author contributions statement

- Conceptualization: NH, TAM, BES, SK, KWL and DRE
- Methodology: NH, SK, TAM, SW, MT, HQ, KL, BES, DRE
- Investigation: NH, SK, TAM, SW, MT, HQ, KL, BD, OH, BES, KWL
- Visualization: NH, KL, BES, DRE
- Funding acquisition: BES, DRE
- Project administration: NH, TAM, BES, DRE
- Supervision: NH, TAM, BES, DRE
- Writing original draft: BES, DRE
- Writing review & editing: NH, SK, TAM, SW, MT, HQ, KL, BD, OH, KWL, BES, DRE

## Supplementary materials

- Supplementary Materials
- Materials and Methods
- Figs. S1 to S8
- Tables S1 to S5

## Notes

### Competing Interest Statement

BES is founder and employee of Specific Biologics Inc. DRE is co-founder of Specific Biologics Inc and is a member of the Scientific Advisory Board. NH, TAM, SK, SW, KL, BD, OH, HQ and MT are employees of Specific Biologics Inc. NH, BES, TAM, DRE are co-inventors on a patent related to the technology presented in this manuscript.

### Summary of Updates

The author list has been updated.

## References

1. Ciccia, A. & Elledge, S. J. The DNA damage response: making it safe to play with knives. Mol. Cell 40, 179–204 (2010).

2. Nambiar, T. S., Baudrier, L., Billon, P. & Ciccia, A. CRISPR-based genome editing through the lens of DNA repair. Mol. Cell 82, 348–388 (2022).

3. Carroll, D. Genome engineering with targetable nucleases. Annu. Rev. Biochem. 83, 409–439 (2014).

4. Kim, H. & Kim, J.-S. A guide to genome engineering with programmable nucleases. Nat. Rev. Genet. 15, 321–334 (2014).

5. Yeh, C. D., Richardson, C. D. & Corn, J. E. Advances in genome editing through control of dna repair pathways. Nat. cell biology 21, 1468–1478 (2019).

6. van Overbeek, M. et al. DNA repair profiling reveals nonrandom outcomes at Cas9-mediated breaks. Mol. Cell 63, 633–646 (2016).

7. Wolfs, J. M. et al. Biasing genome-editing events toward precise length deletions with an RNA-guided TevCas9 dual nuclease. Proc. Natl. Acad. Sci. 113, 14988–14993 (2016).

8. da Silva, J. F. et al. Click editing enables programmable genome writing using DNA polymerases and HUH endonucleases. bioRxiv (2023).

9. Anzalone, A. V. et al. Search-and-replace genome editing without double-strand breaks or donor DNA. Nature 576, 149–157 (2019).

10. Komor, A. C., Kim, Y. B., Packer, M. S., Zuris, J. A. & Liu, D. R. Programmable editing of a target base in genomic dna without double-stranded dna cleavage. Nature 533, 420–424 (2016).

11. Liu, B. et al. Targeted genome editing with a DNA-dependent DNA polymerase and exogenous DNA-containing templates. Nat. Biotechnol. 1–7 (2023).

12. Chen, P. J. et al. Enhanced prime editing systems by manipulating cellular determinants of editing outcomes. Cell 184, 5635–5652 (2021).

13. Li, X. et al. Development of a versatile nuclease prime editor with upgraded precision. Nat. Commun. 14, 305 (2023).

14. Truong, D.-J. J. et al. Exonuclease-enhanced prime editors. Nat. Methods (2024).

15. Schimmel, J. et al. Modulating mutational outcomes and improving precise gene editing at CRISPR-Cas9-induced breaks by chemical inhibition of end-joining pathways. Cell Reports 42 (2023).

16. Leppek, K. et al. Combinatorial optimization of mRNA structure, stability, and translation for RNA-based therapeutics. Nat. Commun. 13, 1536 (2022).

17. Scully, R., Panday, A., Elango, R. & Willis, N. A. DNA double-strand break repair-pathway choice in somatic mammalian cells. Nat. Rev. Mol. cell Biol. 20, 698–714 (2019).

18. Meers, C. et al. Genetic characterization of three distinct mechanisms supporting RNA-driven DNA repair and modification reveals major role of dna polymerase ζ. Mol. Cell 79, 1037–1050 (2020).

19. McDevitt, S., Rusanov, T., Kent, T., Chandramouly, G. & Pomerantz, R. T. How RNA transcripts coordinate DNA recombination and repair. Nat. Commun. 9, 1091 (2018).

20. Storici, F., Bebenek, K., Kunkel, T. A., Gordenin, D. A. & Resnick, M. A. RNA-templated DNA repair. Nature 447, 338–341 (2007).

21. Mazina, O. M., Keskin, H., Hanamshet, K., Storici, F. & Mazin, A. V. Rad52 inverse strand exchange drives RNA-templated DNA double-strand break repair. Mol. Cell 67, 19–29 (2017).

22. Black, S. J. et al. Molecular basis of microhomology-mediated end-joining by purified full-length Polθ. Nat. Commun. 10, 4423 (2019).

23. Carvajal-Garcia, J. et al. Mechanistic basis for microhomology identification and genome scarring by polymerase theta. Proc. Natl. Acad. Sci. 117, 8476–8485 (2020).

24. Chandramouly, G. et al. Polθ reverse transcribes RNA and promotes RNA-templated DNA repair. Sci. Adv. 7, eabf1771 (2021).

25. Chakraborty, A. et al. Human DNA polymerase *η* promotes RNA-templated error-free repair of DNA double-strand breaks. J. Biol. Chem. 299 (2023).

26. Tsegay, P. S. et al. RNA-guided DNA base damage repair via DNA polymerase-mediated nick translation. Nucleic Acids Res. 51, 166–181 (2023).

27. Hamilton, T. A. et al. Efficient inter-species conjugative transfer of a CRISPR nuclease for targeted bacterial killing. Nat. Commun. 10, 4544 (2019).

28. Bell-Pedersen, D., Quirk, S. M., Bryk, M. & Belfort, M. I-TevI, the endonuclease encoded by the mobile td intron, recognizes binding and cleavage domains on its DNA target. Proc. Natl. Acad. Sci. 88, 7719–7723 (1991).

29. Derbyshire, V., Kowalski, J. C., Dansereau, J. T., Hauer, C. R. & Belfort, M. Two-domain structure of the td intron-encoded endonuclease I-TevI correlates with the two-domain configuration of the homing site. J. Mol. Biol. 265, 494–506 (1997).

30. Bryk, M., Belisle, M., Mueller, J. E. & Belfort, M. Selection of a remote cleavage site by i-tevi, thetdintron-encoded endonuclease. J. Mol. Biol. 247, 197–210 (1995).

31. Edgell, D. R., Stanger, M. J. & Belfort, M. Coincidence of cleavage sites of intron endonuclease I-TevI and critical sequences of the host thymidylate synthase gene. J. Mol. Biol. 343, 1231–1241 (2004).

32. Ran, F. A. et al. In vivo genome editing using Staphylococcus aureus Cas9. Nature 520, 186–191 (2015).

33. DeKelver, R. C. et al. Functional genomics, proteomics, and regulatory DNA analysis in isogenic settings using zinc finger nuclease-driven transgenesis into a safe harbor locus in the human genome. Genome Res. 20, 1133–1142 (2010).

34. Kotin, R. M., Linden, R. M. & Berns, K. I. Characterization of a preferred site on human chromosome 19q for integration of adeno-associated virus DNA by non-homologous recombination. The EMBO journal 11, 5071–5078 (1992).

35. Kleinstiver, B. P. et al. Monomeric site-specific nucleases for genome editing. Proc. Natl. Acad. Sci. 109, 8061–8066 (2012).

36. Wolfs, J. M. et al. Megatevs: single-chain dual nucleases for efficient gene disruption. Nucleic Acids Res. 42, 8816–8829 (2014).

37. Kleinstiver, B. P. et al. The I-TevI nuclease and linker domains contribute to the specificity of monomeric TALENs. G3: Genes, Genomes, Genet. 4, 1155–1165 (2014).

38. Strobel, B. et al. A small-molecule-responsive riboswitch enables conditional induction of viral vector-mediated gene expression in mice. ACS Synth. Biol. 9, 1292–1305 (2020).

39. Valley, H. C. et al. Isogenic cell models of cystic fibrosis-causing variants in natively expressing pulmonary epithelial cells. J. Cyst. Fibros. 18, 476–483 (2019).

40. de Serres, F. J. & Blanco, I. Prevalence of α1-antitrypsin deficiency alleles PI* S and PI* Z worldwide and effective screening for each of the five phenotypic classes PI* MS, PI* MZ, PI* SS, PI* SZ, and PI* ZZ: a comprehensive review. Ther. Adv. Respir. Dis. 6, 277–295 (2012).

41. Packer, M. S. et al. Evaluation of cytosine base editing and adenine base editing as a potential treatment for alpha-1 antitrypsin deficiency. Mol. Ther. 30, 1396–1406 (2022).

42. Lombardo, A. et al. Site-specific integration and tailoring of cassette design for sustainable gene transfer. Nat. Methods 8, 861–869 (2011).

43. Ponnienselvan, K. et al. Reducing the inherent auto-inhibitory interaction within the pegRNA enhances prime editing efficiency. Nucleic Acids Res. gkad456 (2023).

44. Srivastava, M. et al. An inhibitor of nonhomologous end-joining abrogates double-strand break repair and impedes cancer progression. Cell 151, 1474–1487 (2012).

45. Huang, F. et al. Identification of specific inhibitors of human rad51 recombinase using high-throughput screening. ACS chemical biology 6, 628–635 (2011).

46. Huang, F. et al. Targeting BRCA1-and BRCA2-deficient cells with RAD52 small molecule inhibitors. Nucleic Acids Res. 44, 4189–4199 (2016).

47. Zatreanu, D. et al. Polθ inhibitors elicit BRCA-gene synthetic lethality and target PARP inhibitor resistance. Nat. Commun. 12, 3636 (2021).

48. Savic, N. et al. Covalent linkage of the DNA repair template to the CRISPR-Cas9 nuclease enhances homology-directed repair. elife 7, e33761 (2018).

49. Li, G., Wang, H., Zhang, X., Wu, Z. & Yang, H. A Cas9–transcription factor fusion protein enhances homology-directed repair efficiency. J. Biol. Chem. 296 (2021).

50. Wyatt, D. W. et al. Essential roles for polymerase θ-mediated end joining in the repair of chromosome breaks. Mol. cell 63, 662–673 (2016).

51. Ramsden, D. A., Carvajal-Garcia, J. & Gupta, G. P. Mechanism, cellular functions and cancer roles of polymerase-theta-mediated dna end joining. Nat. Rev. Mol. Cell Biol. 23, 125–140 (2022).

52. Schimmel, J., Kool, H., van Schendel, R. & Tijsterman, M. Mutational signatures of non-homologous and polymerase theta-mediated end-joining in embryonic stem cells. The EMBO journal 36, 3634–3649 (2017).

53. Zimmerly, S., Guo, H., Perlman, P. S. & Lambowltz, A. M. Group II intron mobility occurs by target DNA-primed reverse transcription. Cell 82, 545–554 (1995).

54. Luan, D. D., Korman, M. H., Jakubczak, J. L. & Eickbush, T. H. Reverse transcription of R2Bm RNA is primed by a nick at the chromosomal target site: a mechanism for non-LTR retrotransposition. Cell 72, 595–605 (1993).

55. Zhang, X. et al. Harnessing eukaryotic retroelement proteins for transgene insertion into human safe-harbor loci. Nat. Biotechnol. (2024).

56. Mueller, J. E., Smith, D., Bryk, M. & Belfort, M. Intron-encoded endonuclease I-TevI binds as a monomer to effect sequential cleavage via conformational changes in the td homing site. The EMBO J. 14, 5724–5735 (1995).

57. Kleinstiver, B. P., Wolfs, J. M. & Edgell, D. R. The monomeric GIY-YIG homing endonu-clease I-BmoI uses a molecular anchor and a flexible tether to sequentially nick DNA. Nucleic Acids Res. 41, 5413–5427 (2013).

58. Van Roey, P., Meehan, L., Kowalski, J. C., Belfort, M. & Derbyshire, V. Catalytic domain structure and hypothesis for function of GIY-YIG intron endonuclease I-TevI. Nat. Struct. Biol. 9, 806–811 (2002).

59. Roy, A. C., Wilson, G. G. & Edgell, D. R. Perpetuating the homing endonuclease life cycle: identification of mutations that modulate and change I-TevI cleavage preference. Nucleic Acids Res. 44, 7350–7359 (2016).

60. Edgell, D. R., Stanger, M. J. & Belfort, M. Importance of a single base pair for discrimination between intron-containing and intronless alleles by endonuclease I-BmoI. Curr. Biol. 13, 973–978 (2003).

61. Edgell, D. R. & Shub, D. A. Related homing endonucleases I-BmoI and I-TevI use different strategies to cleave homologous recognition sites. Proc. Natl. Acad. Sci. 98, 7898–7903 (2001).

