## Supplementary data for "Harnessing RNA-based DNA repair pathways for targeted gene editing"

**The PDF file includes:**

Materials and Methods
Supplementary Text
Figs. S1 to S9
Tables S1 to S6

### Materials and Methods

#### Cell lines and culturing

All cell lines were cultured at 37°C and 5% $CO_2$. Human embryonic kidney cells for transfection (HEK293, ATCC CRL-1573) or viral particle production (HEK293T; Takara Bio 632273) were cultured in DMEM (Sigma Aldrich D5796-500ML) containing 10% Fetal Bovine Serum (Sigma Aldrich F1051-100mL) and streptomycin and penicillin (Sigma Aldrich P4333-100mL). Engineered monoclonal human bronchial epithelial cells (16HBEge) that were gene edited to contain the *CFTR* DF508, G542X or W1282X mutations were a gift of the Cystic Fibrosis Foundation and cultured as described previously (*1*). Untransformed liver fibroblasts containing the *SERPINA1* c.1096G>A (p.Glu342Lys) mutation (GM11423; Corriell Research Institute) were cultured per manufacturer's directions.

#### Cloning and plasmid construction

For parental plasmids, fragments carrying MALAT, tRNA-ribozyme, and sgRNA cassette were obtained from Integrated DNA Technologies and cloned into pX458 (*2*) and pAAV plasmids carrying relevant nucleases using Gibson assembly. To generate plasmids with relevant rep-gRNAs, fragments carrying modified tRNA-ribozyme and target sequence were assembled with fragments carrying the gRNA scaffold and repair template. The assembled product was inserted into the plasmid backbone via Gibson assembly. All fragments were generated using the polymerase chain reaction (PCR) with GXL PrimeSTAR DNA polymerase (Takara Bio R050). For cloning of gRNAs, oligos pairs were annealed inserted using Golden-Gate into parental plasmids carrying empty gRNA cassettes using established protocol (*3*). Constructs in pX458 and pAAV plasmids were transformed into Top10 (NEB C3019) and Stable (NEB C3040) competent cells, respectively. Correct constructs were identified using PCR screens and verified by Sanger sequencing (The Hospital for Sick Children, Toronto, ON) and whole plasmid sequencing (Plasmidsaurus, Eugene, OR). A list of oligonucleotides used in this study is found in Table S2.

#### Rep-gRNAs

*AAVS1* targeting rep-gRNAs were transcribed *in vitro* using the HiScribe T7 High Yield RNA Synthesis Kit (NEB E2040) from PCR amplicons substrates. Rep-gRNAs targeted to the *CFTR* DF508, G542X and W1282X or *SERPINA1* E342K sites were purchased with 2`-O-methyl-phosphorothioiate modifications on the first three ribonucleotides at the 5'-end from Agilent or Integrated DNA Technologies. A list of rep-gRNA sequences is in Table S1 and oligonucleotides are in Table S2.

#### Protein purification

Plasmids for expressing C-terminal 6X histidine-tagged TevSaCas9 or SaCas9 were transformed into *Escherichia coli* T7 express (NEB C2566), grown in 1L of 2X-YT media (Bio Basic SD7019) in the absence of glucose to an $A_{600}$ of 0.6 – 0.8 and expression was induced using 1 mM IPTG (Bioshop IPT001.10). Cells were harvested by centrifugation at 12,000 x g and lysed with sonication (Banson) at 30% amplitude for 7 minutes using 10 second on/off pulses on ice or with high pressure homogenization with a single pass at 16 – 18,000 psi using an Avestin C5 in lysis buffer (50 mM Tris-Cl pH 8, 10 mM imidazole, 500 mM sodium chloride and 5% glycerol). The lysate was centrifuged for 30 minutes at 14,000 x g and the supernatant applied to a nickel-charged 5 mL His-trap column (Cytiva) equilibrated in lysis buffer. The column was washed with 20 column volumes of lysis buffer and eluted with a 10-column volume linear gradient of lysis buffer to elution buffer (lysis buffer with 500 mM imidazole) in 5 mL fractions. TevSaCas9- or SaCas9-containing fractions were pooled and exchanged into storage buffer (50 mM Tris-Cl pH 8, 250 mM sodium chloride, 5% glycerol and 1 mM DTT) or nucleofection buffer (50 mM Tri-Cl pH 7.4, 300 mM sodium chloride, 0.1 mM EDTA, 50% glycerol and 1 mM DTT) using at least 3 volume exchanges on a 100 kDa MWCO filter (Cytiva).

**Messenger RNA synthesis**

To generate messenger RNA (mRNA), transcription templates carrying a T7 promoter were generated via PCR or maxiprep of the relevant constructs. For maxiprep plasmid templates, DNA was precipitated with 100% pre-chilled isopropanol at the ratio of 10:7 (v:v) for 20 minutes on ice, followed by 4 times washes in fresh pre-chilled 70% Ethanol. PCR-based templates were purified using GenepHlow PCR clean-up kit (Geneaid DFH300). Purified templates were tested for traces of RNAse using RNAse Alert test kit (Invitrogen AM1964 or IDT 11-04-03-03/11-04-02-07). mRNA was transcribed *in vitro* from 1 mg DNA template input using the HiScribe T7 High Yield RNA Synthesis Kit (NEB E2040) with or without a 3-O-Me-m7G(5')ppp(5')G RNA 5' Cap Structure Analog [C3] (NEB S1411) or CleanCap AG [C1] (TriLink N-7113). For capped reactions, a molar ratio of 6:5:5:1:4 of ATP:UTP:CTP:GTP:Cap was used for the C3 cap or 6:5:5:5:4 for the C1 cap. To minimize degradation in the presence of trace RNAse, 40 units of Murine RNAse inhibitor (NEB M0314) was added per 20 µL reaction. *In vitro* transcription was carried out for 120 minutes at 32°C, followed by poly A tailing reaction by incubating mRNA samples with 15 units of Poly A polymerase (NEB M0276) and 4 units of DNase I (NEB M0303) in 50 µL reaction for 60 minutes at 32°C. Reactions were then purified using a Monarch RNA purification kit (New England Biolabs) following manufacturer's directions except 100% ethanol was replaced with 100% isopropanol. RNAs were quantified using Qubit RNA broad range kit (Invitrogen Q10210), RNA integrity was evaluated using Qubit RNA IQ kit (Invitrogen Q33221). In addition, capping efficiency was evaluated via Ribozyme based cleavage assay and the presence of intact regions (e.g. I-TevI, Cas9, rep-gRNA, etc.) of the mRNAs were evaluated using RT-PCR. RNAs are listed in Table S3.

**Adeno-associated virus (AAV) production**

HEK293T cells were transfected with the AAV2 Helper Free Packaging System (Cell Biolabs Inc) and pAAV-TevCas9 or pAAV-SaCas9 vector. Cell supernatant was collected, and viral genome copy number was measured by quantitative PCR. AAVs used in this study are listed in Table S4.

**HEK293 and 16HBEge transfections**

HEK293 cells were seeded in TC-treated 48-well plates (Corning CLS3548) at a density of 60,000 cells per $cm^2$ approximately 24 hours prior to transfection in DMEM without antibiotics. For plasmid DNA, 500 ng was transfected using 0.5 µL of 1 mg/ml linear polyethyleneimine, MW 25,000 (Alfa Aesar AA4389601) for 24-48 hours before cells were harvested. For all-in-one mRNA transfections, 500 ng of mRNA were transfected with Trans-IT X2 reagent (Mirus) using the manufacturer's protocol and cells harvested 24 hours after transfection. For mRNA with rep-gRNA co-transfections 50 nM of rep-gRNA was also included. For ribonucleoprotein complex transfection, purified TevSaCas9 or SaCas9 were incubated at 37°C with rep-gRNA or gRNA for 10 minutes to form RNP complex. Complexes were transfected using Trans-IT X2 reagent (Mirus) using the manufacturer's protocol and cells harvested 24 hours after transfection. For 16HBEge transfections, cell culturing flasks and plates were coated for 2-3 hours at 37°C with a solution of 1% (v/v) bovine serum albumin faction V (Gibco 15260037), 30 mg/mL Bovine collagen solution, Type 1 (Advanced BioMatrix,Inc 5005) and 10 mg/mL Fibronectin from human plasma (ThermoFisher Scientific 33016015). 16HBEge were seeded in coated 48-well plates (Corning CLS3548) at a density of 17,500 cells per $cm^2$ approximately 24 hours before transfection in Minimum Essential Medium (Gibco 11095080) with 1X Penicillin-Streptomycin (100X Millipore-Sigma P4333). Prior to transfection, media was substituted for antibiotic free media.

**Nucleofection**

16HBEge cells were grown overnight to 60–70% confluency in treated T-flasks. Cells were harvested and resuspended in Buffer R for nucleofection using a Neon nucleofector following manufacturers protocol (Invitrogen). For ribonucleoprotein complex nucleofections, 1,500,000 cells were nucleofected using a 100 µL tip with 1.2 µM of purified TevSaCa9 or SaCas9 protein complexed for 15 minutes at room temperature with 1.0 µM of rep-gRNA or sgRNA. Cells were nucleofected with 1650 V for 10 milliseconds with 2 pulses. Nucleofected cells were grown in antibiotic-free media in 24-well plates for 24 hours before harvesting for genotyping. GM11423 cells were grown overnight to 70–90% confluency in treated T-flasks. Cells were harvested and resuspended in Buffer R for nucleofection using a Neon nucleofector following manufacturers protocol (Invitrogen). 100,000 cells were nucleofected using a 10 µL tip with 2 µg of TevSaCa9 *SERPINA1* E342K mRNA without and with 200 nM of additional *SERPINA1* E342K rep-gRNA. Cells were nucleofected with 1150 V for 20 milliseconds with 2 pulses. Nucleofected cells were grown in antibiotic-free media in 24-well plates for 24 hours before harvesting for genotyping.

**Transduction**

HEK293 cells were seeded in TC-treated 48-well plates (Corning CLS3548) at a density of 50,000 cells per $cm^2$ approximately 24 hours prior to transduction in Dulbecco's Modified Eagle's Medium - high glucose (Millipore-Sigma D5796). Cells were transduced at an MOI of 1,000 to 10,000, centrifuged at 800 x g for 1 hour at 37°C and incubated at 37°C and 5% $CO_2$ for 24 hours. After 24 hours, virus was inactivated with 4% (v/v) formaldehyde for 10 minutes and dissociated with 1.25% (v/v) of 0.5M EDTA, pH 8.0, centrifuged 2,000 x g and washed and centrifuged three times with ice cold phosphate buffered saline pH 7.4 at 2,000 x g before final centrifuged at 2,000 x g.

**Inhibition of repair pathways**

B02 (Sigma Aldrich SML0364) was used to inhibit HDR at three doses of 1.0 µM, 10.0 µM and 100 µM (*4*, *5*). SCR7 (Sigma Aldrich SML1546) was used to block NHEJ at final concentrations of 0.01 µM, 0.1 µM, and 1.0 µM (*6*). D-I03 (MedChem Express HY-124691) was used to inhibit Rad52 at the range of 2.5 nM, 25 nM, and 250 nM (*7*). Polq inhibition was done using ART558 (MedChem Express HY-141520) at final concentrations of 0.01 µM, 0.1 µM, 1.0 µM (*8*). HEK293 cells were seeded in TC-treated 48-well plates (Corning CLS3548) at a density of 60,000 cells per $cm^2$

approximately 18-24 hours prior to transfection in DMEM. Four hours before transfection or transduction, inhibitors were added at above-mentioned concentrations.

**Genomic DNA extraction and amplicon generation**

Harvested cells were resuspended to approximately 40 μL in QuickExtract (Lucigen LGN-QE09050) per 100,000 cells and genomic DNA extracted per manufacturer's protocol. PCR amplicons were generated using 1–3 μL of genomic extract with PrimeStar GXL polymerase (Takara Bio R050) using standard reaction conditions and the primers in Table S1.

**Restriction enzyme digests for editing and repair analysis**

Approximately 100–200 ng of each target site PCR amplicon was digested for 30 minutes at 37°C from treated or mock treated samples with the appropriate restriction enzyme. BglII (ThermoFisher Scientific ER0081) was used for *AAVS1* amplicons, SspI (NEB R3132) for *CFTR* DF508 and G542X amplicons, HindIII (NEB R3104) for *CFTR* W1282X, and MfeI (NEB R3589) for *SERPINA1* E342K. After heat inactivation, the reaction was resolved on a 1.5% Tris Borate EDTA (TBE) agarose gel and the density of digested and undigested bands was quantified using ImageJ as a function of total DNA.

**Western blot analysis**

To evaluate endogenous protein expression, cells were either lysed using Triprep kit (Takara Bio 740966.50) or RIPA lysis buffer (NEB 9806S) with 1x proteinase inhibitor (Roche 11873580001). For cell lysis using RIPA lysis buffer, buffer was warmed to 50°C to dissolve all precipitate and added to cells at the ratio of 100 μL per 1,000,000 cells and mixed by pipetting up and down 5-10 times. Lysis was done on ice with 200-220 rpm shaking for 10 minutes, followed by centrifugation at 16,000 x g for 10 minutes at 40°C. Cell lysate was quantified using Qubit protein assay broad range (Invitrogen A50668) or Nanodrop. 10-20 μg of cell lysate was loaded into 4-20% Tris-Glycine gel (Invitrogen XP04120BOX). For co-immunoprecipitation (coIP), HEK293 cells expressing nucleases and rep-gRNAs were lysed using NP40 cell lysis buffer. Prior to coIP, HA-agarose bead (GenScript L00777) was equilibrated to room temperature for 15 minutes. Per sample, 20 μL of beads were washed with NP40 lysis buffer for 4 times before being combined with lysate and incubate on rocking shaker for 1 hour at 40°C. Beads were then collected and washed in pre-chilled PBS for 4 times by centrifugation at 8,000 x g for 10 seconds and eluted in SDS-based loading buffer. The SDS-PAGE gel for western blotting was run at 150V for 45-60 minutes and transferred using the Trans-Blot turbo transfer kit (BioRad 1704273) at 25V, 2.5A, for 8-9 minutes. The membrane was blocked and immunostained using BSA-based buffer. Detection was done using Clarity Western ECL Substrate (BioRad 1705061) and images were captured using ChemiDoc Imaging System (BioRad). Antibody was used at the final ratio of 1:1,000 for Beta-Actin antibody HRP (Invitrogen PA1-183-HRP), 1:1,000 for anti-Rad52 (Invitrogen PA5-119963), 1:1,000 for Polq antibody (Invitrogen PA5-115130), 1:1,000 for anti-HA HRP (Invitrogen 26183-HRP), 1:20,000 for Goat anti-mouse HRP (Invitrogen 31430), and 1:20,000 for Goat anti-rabbit HRP (Invitrogen 31460).

**Quantitative reverse transcription polymerase chain reaction (RT-qPCR)**

RNA was extracted using the Triprep Kit (Takara Bio 740966.50). Samples were centrifuged at 8,000 g for 8 sec between steps and at 15,000 x g for 3 minutes at the final elution. Quantification was done using Qubit or Nanodrop using elution buffer as reference. 1 μg of extracted RNA was used to make cDNA using LunarScript RT Supermix (NEB E3010) following manufacturer's protocol. Generated cDNAs were then used for RT-PCR using relevant oligonucleotides (Table S1) and Luna Universal qPCR master mix (NEB M3003). Reactions were carried out using QuantStudio 6 pro following previously established conditions (*9*). Data was analyzed using ddCT method for differential expression.

**RNA processing assay**

To evaluate *trans*-cleavage efficiency of the ribozyme-tRNA in the all-in-one construct, mRNA was synthesized and purified as described above. HEK293 cells were lysed in NP40 cell lysis buffer (Invitrogen FNN0021). Genomic DNA and total RNA were removed from lysate using 3 rounds of Triprep kit, leaving the protein in the flow through. Protein supernatants were dialyzed against 1x PBS. For the RNA processing assay, 250 ng of mRNA was incubated with 10 μg of protein supernatant at 37°C for desired periods of time. To stop cleavage reaction, proteinase was added at the final concentration of 0.05 μg/mL and incubated at 37°C for 30 minutes and run on 1.5% TBE gel at 120V for 75 minutes, stained with Ethidium Bromide for 10 minutes and destained in deionized water for 20 minutes before visualization.

**Next generation sequencing and analysis**

Target site amplicons were sequenced by GeneWiz by Azenta Life Sciences (New Jersey) or The Centre for Applied Genomics (Toronto, Ontario) for 250-bp paired end Illumina sequencing. Analysis of editing was performed using CRIS.py (*10*) or CRISPresso2 (*11*). For CRIS.py analysis, individual R1 and R2 fastq file were first merged using FLASH (*12*). Editing was assessed using the available CRIPS.y2 python script modified to contain the unedited sequence (ref_seq) and the expected edited sequence (test_seq) for each target site excluding the 20-bp corresponding to amplicon specific PCR primers. For CRISPResso2 analysis, individual R1 and R2 fastq files were used to compare editing using the expected (--expected_hdr_amplicon_seq) and unedited reference sequence (--amplicon_seq) using the gRNA portion of the rep-gRNA as the input for –guide_seq. For analysis of predicted off-target sites for the *AAVS1* rep-gRNA, we used CRIPRessoPooled with a file of off-target amplicon sequences (Table S4). Fastq files were further separated by delivery method for off-target analysis (6 replicates each for AAV, RNP, pDNA, and mRNA delivery) and editing reported as the mean value for all replicates where each off-target site had at least 10000 reads (--min_reads_to_use_region 10000). For the rep-RNA deletion experiment, read length relative to the unmodified length for edited or mock treated cells was determined using custom Perl and R scripts.

**Off-target analysis**

SaCas9-dependent off-target sites with 4 or less mismatches and no RNA or DNA bulges for the *AAVS1* gRNA binding site in the GRCh38/hg38 human genome were identified using Cas-OFFinder (*13*) (Table S5). The gRNA site, PAM sequence, and 35-bp upstream region were extracted from GRCh38/hg38 human sequence using the R package Bsgenome. The genomic coordinates of the on- and off-target sites were used to design a multiplex PCR primer pool using the rhAmpSeq design tool (Integrated DNA Technologies). Amplicons were generated using the rhAmpSeq protocol and sequenced using a paired-end Illumina MiSeq v2 500-cycle run (The Centre for Applied Genomics, Toronto, Ontario). Percent editing was analyzed using CRISPAltRations (Integrated DNA Technologies) or CRISPECTOR.

**In vitro cleavage assays**

1 μM of purified TevSaCa9 or SaCas9 protein complexed for 15 minutes at room temperature with 1 μM of rep-gRNA or sgRNA. 100 nM of RNP complexes were added to 10 nM of substrate in 1X New England Biolabs Buffer 2 at 37°C to start the reaction. After the indicated time, 20 μL of reactions were removed and stopped with 10 μL of 50 mM EDTA, pH 8.0, 150 μg / mL RNase A and 300 μg / mL Proteinase K for 30 minutes for 30 minutes at 37°C. 6 μL of 6X Loading Dye was added to each stopped reaction and half the reaction was run on 1.5% TBE gel at 120V for 75 minutes, stained with Ethidium Bromide for 10 minutes and destained in deionized water for 20 minutes before imaging.

### Supplementary Text

#### Editing outcomes at the *CFTR* G452X target site

With TevSaCas9/rep-gRNA, we observed 51 +/- 8% editing at the *CFTR* G542X site (Fig. SDE). To compare editing outcomes relative to SaCas9 with a standard sgRNA approach targeted to the same site, we transfected 16HBEge cells with RNPs consisting of SaCas9/rep-gRNA or SaCas9 with a sgRNA and a single-stranded DNA donor template (ssODN) that requires HDR pathways for repair. With SaCas9/rep-gRNA, 99% of deep sequencing reads corresponded to unmodified sites, with <1% of reads consistent with a-EJ repair (Fig. S4H and S4J). With DNA substrates *in vitro*, we observed robust cleavage with SaCas9/rep-gRNA RNPs, suggesting that the lack of observed activity in cells is not due to an inactive RNP (Fig. S1C). In contrast, co-delivery of SaCas9 with an sgRNA and a ssODN resulted in the characteristic spectrum of indels observed with Cas9 editing that are consistent with a-EJ repair (27% of reads) (Fig. S5H and S5K). Less than 1% of reads were consistent with correct HDR repair (Fig. S5H). These data show that rep-editing outcomes are much more uniform that with a standard sgRNA or ssODN repair template at the same target site. Moreover, the data indicate that editing with SaCas9 and a rep-gRNA does not support DNA repair in cells.

#### Specificity of rep-editing

The mismatch data in Fig. 3 and our previous characterization of TevCas9 fusions demonstrates that the gRNA and PAM are the primary targeting determinants (*14*). Any off-target cleavage is derived from binding and cleavage to mismatched SaCas9/gRNA sites rather than from independent binding and cleavage by the Tev nuclease domain at CNNNG motifs. We assessed in greater depth off-target cleavage by TevSaCas9/*AAVS1* rep-gRNAs by computationally identifying 38 off-target sites to the *AAVS1* site (Fig. S7) (*13*). These included 6 sites with 3 mismatches and 32 sites with 4 mismatches to the *AAVS1* gRNA portion of the rep-gRNA (there were no identified off-target sites with 1 or 2 mismatches; Fig. S7). Examination of sequence upstream of each of the off-target gRNA binding sites revealed multiple CNNNG motifs. Only 3 off-target sites (each with 4 gRNA mismatches, marked with an asterisk) contained CAGGG motifs that would support *AAVS1*/rep-gRNA-mediated repair (Fig. S7). We assessed off-target editing by independently using AAV or mRNA delivery of TevSaCas9/*AAVS1* rep-gRNA to HEK293 cells and by using a multiplex PCR approach with a pool of primers designed against the on- and off-target sites followed by deep sequencing. We found that on-target editing ranged between 34-80% (Fig. S8). Using established off-target analysis methods of analysis (*15, 16*), we did not observe significant levels of cleavage at the 38 off-target sites. For instance, off-target sites OT_3 and OT_30 showed very low levels of editing (less than 0.2% by CRISPECTOR analysis) but all indels mapped to the SaCas9 cut site and not to the Tev cut site (Fig. S8). We also did not detect any translocations between the on- or off-target sites by CRISPECTOR analysis.

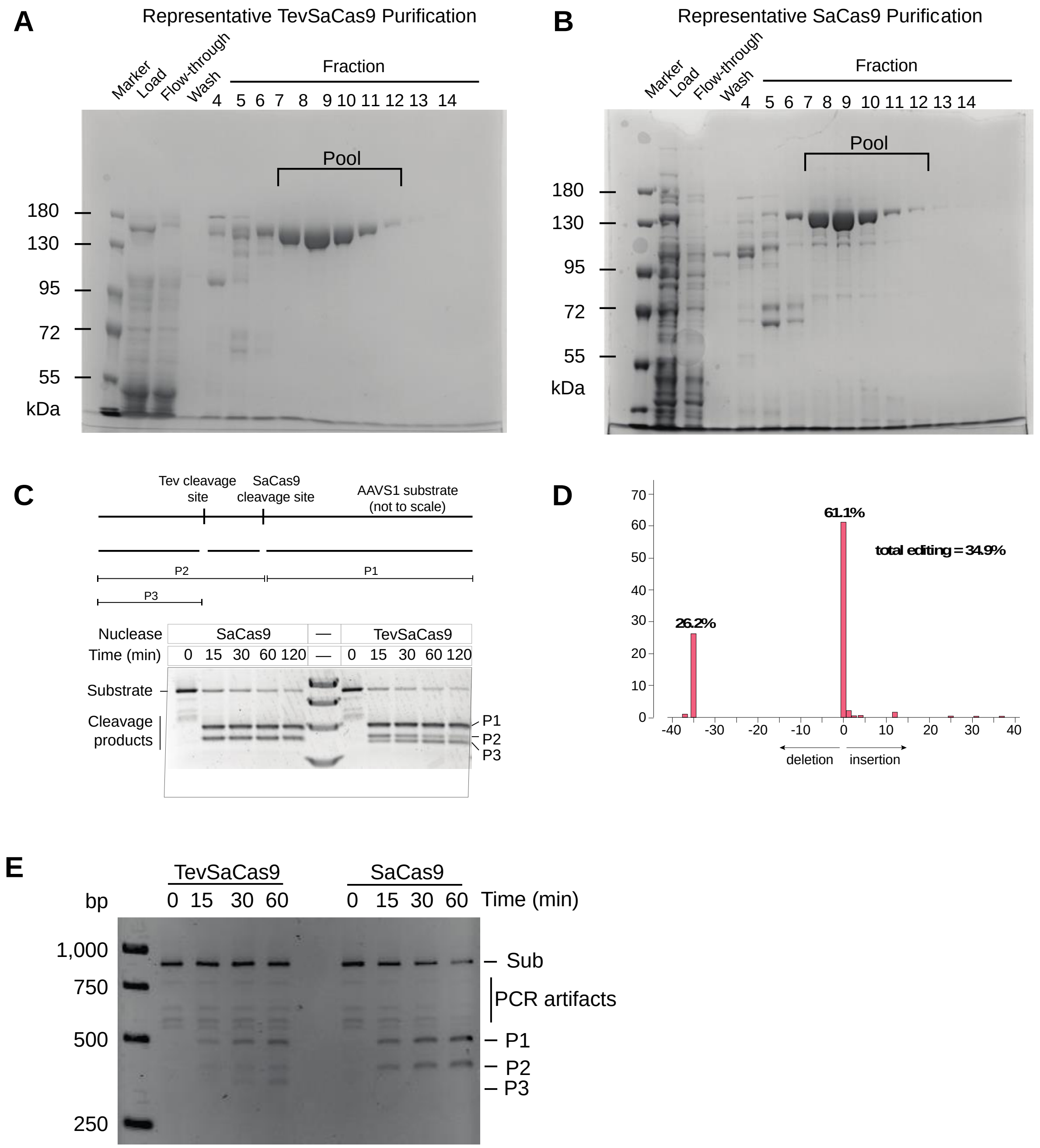


**Fig. S1. Purification and activity of TevSaCas9 and SaCas9. A** and **B**, Representative purifications of TevSaCas9 and SaCas9. Shown are 8% SDS-polyacrylamide gels with fractions indicated above the gels. **C**, Activity of TevSaCas9 and SaCas9 programmed with an *AAVS1* targeting rep-gRNA on a PCR generated *AAVS1* substrate. The predicted cleavage products are indicated above the gel. Reactions were incubated for the indicated time, stopped, and analyzed on a 1.5% agarose gel. **D**, TIDE analysis (*17*) of TevSaCas9 editing of the *AAVS1* site in HEK293 cells.

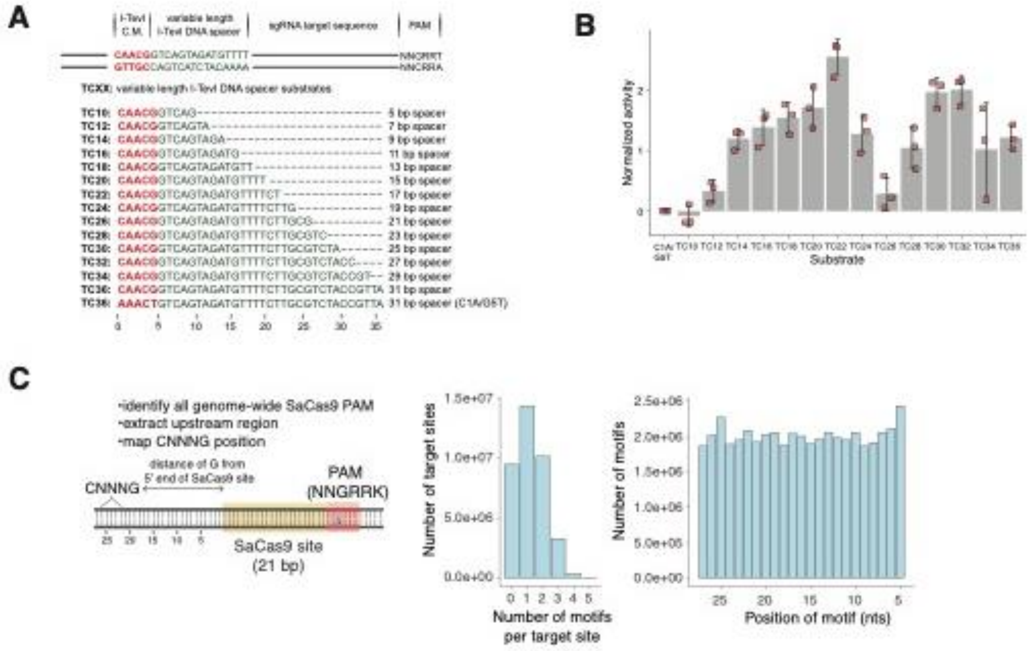


**Fig. S2. Impact of CNNNG motif spacing on TevSaCas9 cleavage. A**, Design of TCXX substrates with the CNNNG motif (CAACG) spaced from 5 to xx bps from the gRNA binding site. Substrates are designated by the distance of the G of the motif from the 5’ end of the gRNA binding site. **B**, Cleavage activity of TevSaCas9 on the TCXX substrates normalized to activity on a substrate with a CNNNG knockout (AAACC). Bars are mean activity of three biological replicates with error reported as standard deviation from the mean. Each dot represents an independent replicate. **C**, Distribution of CNNNG motifs and SaCas9 sites in the human genome.

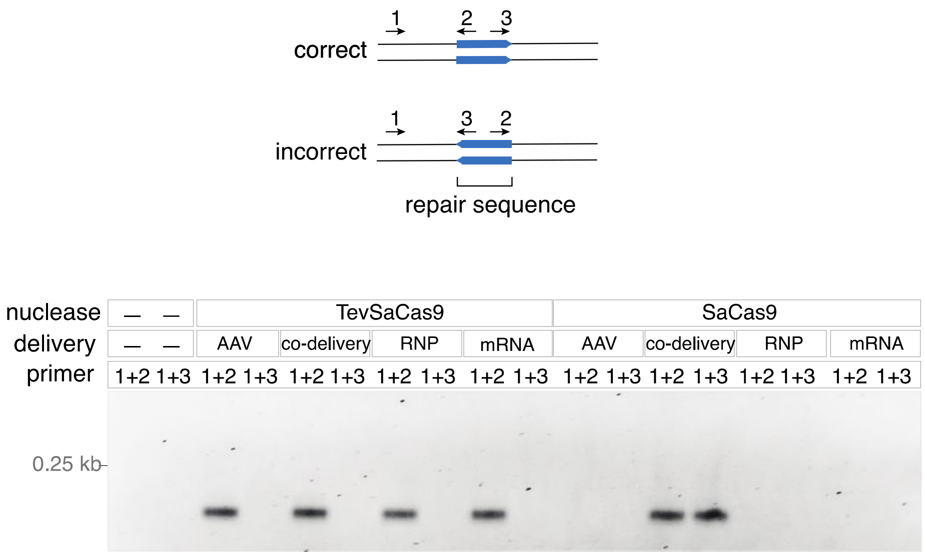


AAV: all-in-one (nuclease + rep-gRNA)
co-delivery: nuclease + gRNA + repair template (ssODN)
RNP: nuclease + rep-gRNA
mRNA: all-in-one (nuclease + rep-gRNA)

**Fig. S3. PCR screen for directional repair at the *AAVS1*-T1 site.** Top, schematic of PCR primers used to identify correctly oriented repair versus incorrectly oriented repair at the *AAVS1* genomic locus. Bottom, agarose gel of diagnostic PCR from total DNA isolated from HEK293 cells treated with the indicated constructs and primer pairs.

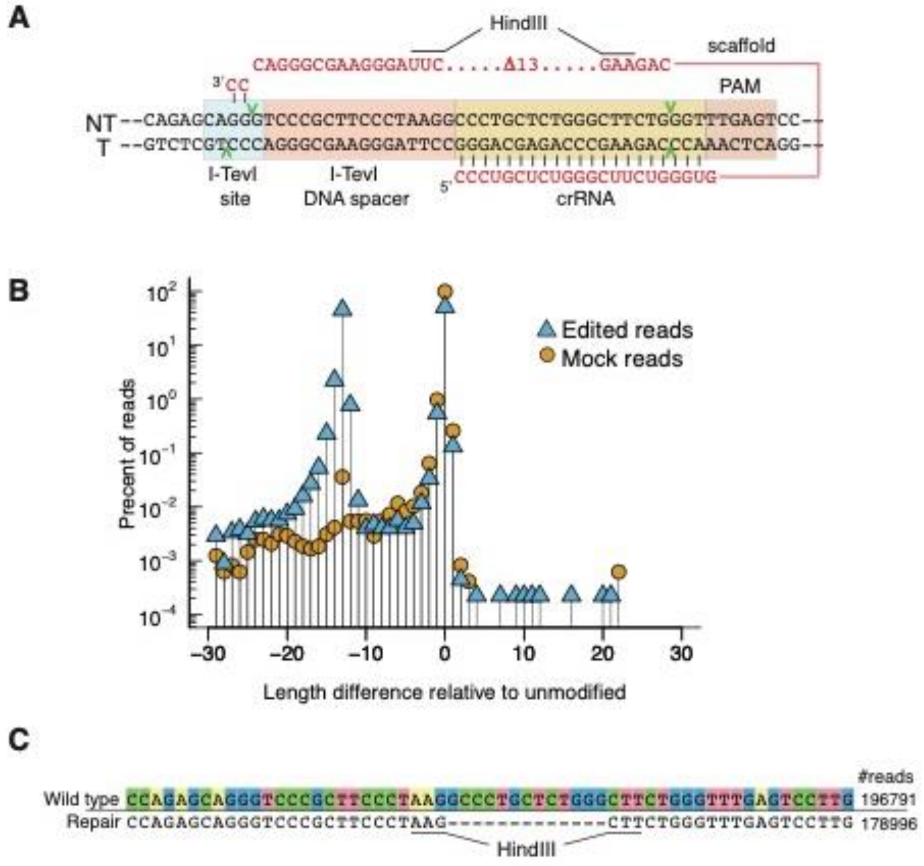


**Fig. S4. Using rep-editing to make a targeted deletion. A,** Design of a rep-gRNA to introduce a 13-bp deletion in the *AAVS1* safe harbor site, with individual components of the rep-gRNA labeled as in Fig. 1B. **B,** Plot of length difference relative to the length of the unmodified *AAVS1* site for deep sequencing reads of HEK293 cells transfected with TevSaCas9/rep-gRNA or mock transfected. **C,** The most prevalent deep sequencing reads from HEK293 cells transfected with TevSaCas9/rep-gRNA. Deletions are indicated by a dash (-) while identities are not colored.

### Figure S5

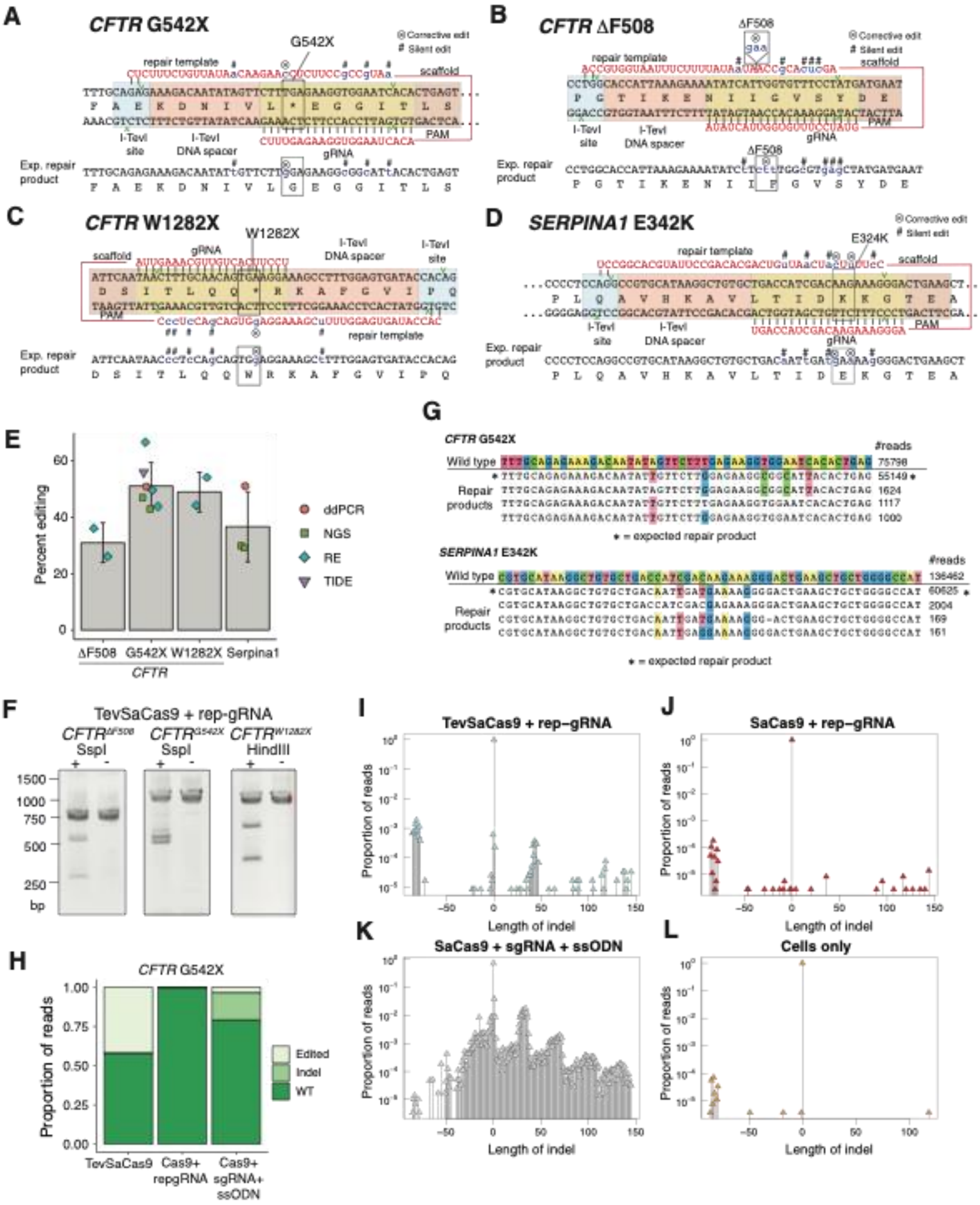
A
CFTR G542X
G542X
Corrective edit
Silent edit
repair template
scaffold
I-TevI site
I-TevI DNA spacer
gRNA
PAM
Exp. repair product
B
CFTR ΔF508
ΔF508
Corrective edit
Silent edit
repair template
scaffold
I-TevI site
I-TevI DNA spacer
gRNA
PAM
Exp. repair product
C
CFTR W1282X
W1282X
scaffold
gRNA
I-TevI DNA spacer
I-TevI site
PAM
repair template
Exp. repair product
D
SERPINA1 E342K
E324K
Corrective edit
Silent edit
repair template
scaffold
I-TevI site
I-TevI DNA spacer
gRNA
PAM
Exp. repair product
E
Percent editing
ddPCR
NGS
RE
TIDE
ΔF508
G542X
W1282X
Serpina1
CFTR
G
CFTR G542X
#reads
Wild type
Repair products
* = expected repair product
SERPINA1 E342K
#reads
Wild type
Repair products
* = expected repair product
F
TevSaCas9 + rep-gRNA
SspI
SspI
HindIII
bp
H
CFTR G542X
Proportion of reads
Edited
Indel
WT
TevSaCas9
Cas9+ repgRNA
Cas9+ sgRNA+ ssODN
I
TevSaCas9 + rep−gRNA
Proportion of reads
Length of indel
J
SaCas9 + rep−gRNA
Proportion of reads
Length of indel
K
SaCas9 + sgRNA + ssODN
Proportion of reads
Length of indel
L
Cells only
Proportion of reads
Length of indel

**Fig. S5. Rep-editing at therapeutically relevant target sites. A-D**, Rep-gRNA design for targeting of the G542X (A), DF508 mutation (B) or W1282X (C) mutations of the *CFTR* gene or E342X mutation in the *SERPINA1* gene. The expected edited product is shown below, with silent substitutions indicated by a hash symbol (#) and corrective edits indicated by a x-marked circle (⊗ ). **E**, Summary of percent editing at the *CFTR* G542X, DF508 or W1282X sites in 16HBEge cells or the *SERPINA1 E342K* site in GM11423 primary liver fibroblast cells. Bars are the mean of editing assessed by the indicated methodology, with bars representing standard deviation from the mean. NGS, next generation sequencing; ddPCR, digital droplet PCR; RE, restriction enzyme digest; TIDE, deconvolution of Sanger sequencing reads. **F** Representative gel of SspI (DF508 or G542X) or HindIII (W1282X) restriction digestion analysis of PCR amplicons of 16HBEge cells. TevSaCas9/rep-gRNA treated cells are indicated by a (+) and transfection reagent only treated cells with a (-) **H**, Summary of NGS data indicating editing outcomes at the G542X site for the indicated treatments. **D**, Alignment of prevalent reads from NGS of PCR products amplified from total 16HBEge *CFTR* G542X or GM11423 *SERPINA1* E342K cells transfected with TevSaCas9/rep-gRNA (cells were not enriched or selected prior to analysis). **I-L,** Plots of the proportion of NGS reads versus to the length difference relative to the unmodified site from 16HBEge cells transfected with the indicated constructs.

A

B

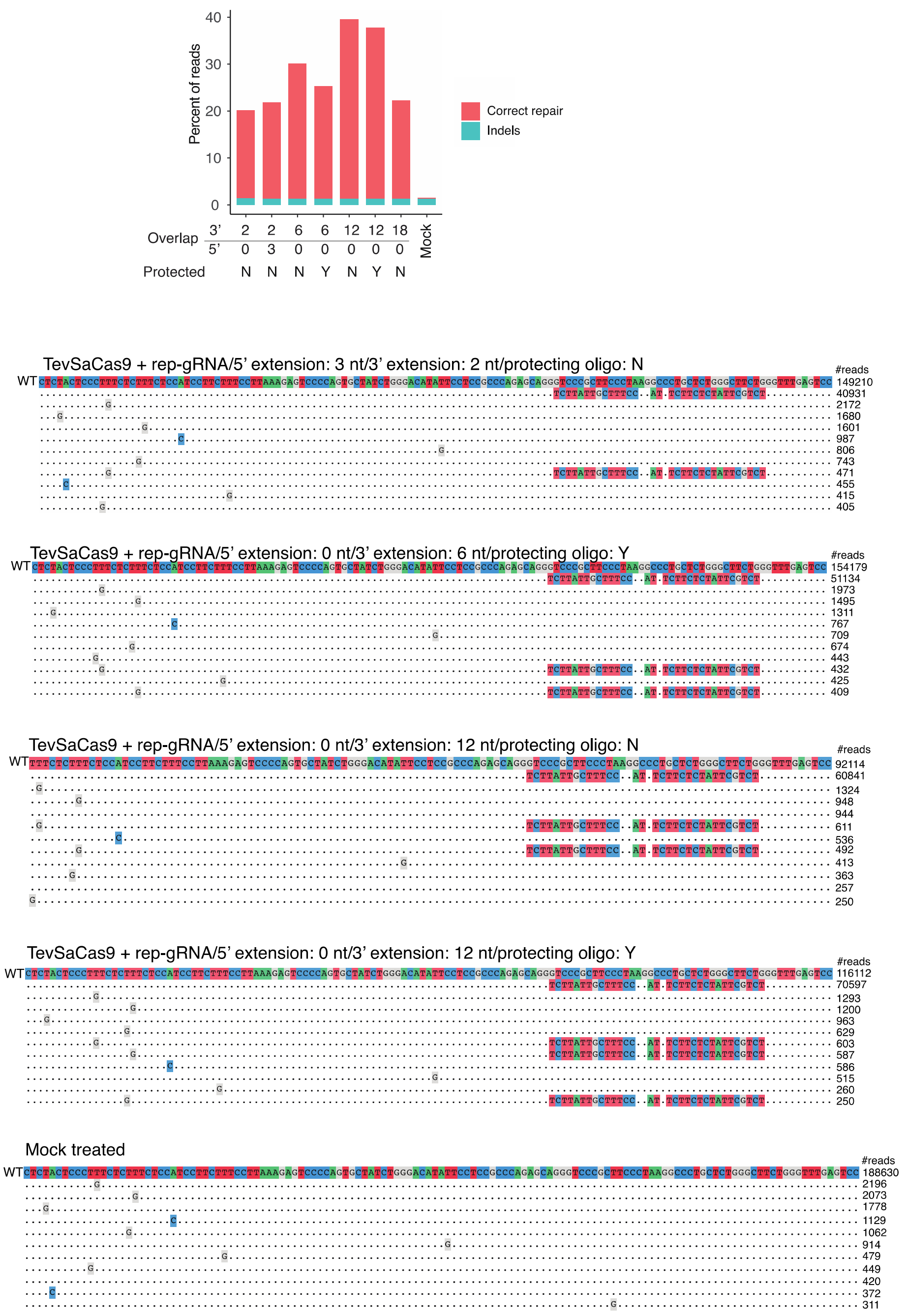


**Fig. S6. Deep sequencing analysis of HEK293 cells transfected with TevSaCas9 and rep-gRNA with different 3' overlaps. A**, Plot of percent editing at the *AAVS1* site with rep-gRNAs containing the indicated 3' or 5' overlaps. **B,** Example deep sequencing reads for the indicated experiments. Identical nucleotides are indicated by dots and nucleotide differences are colored. The number of reads are indicated on the right.

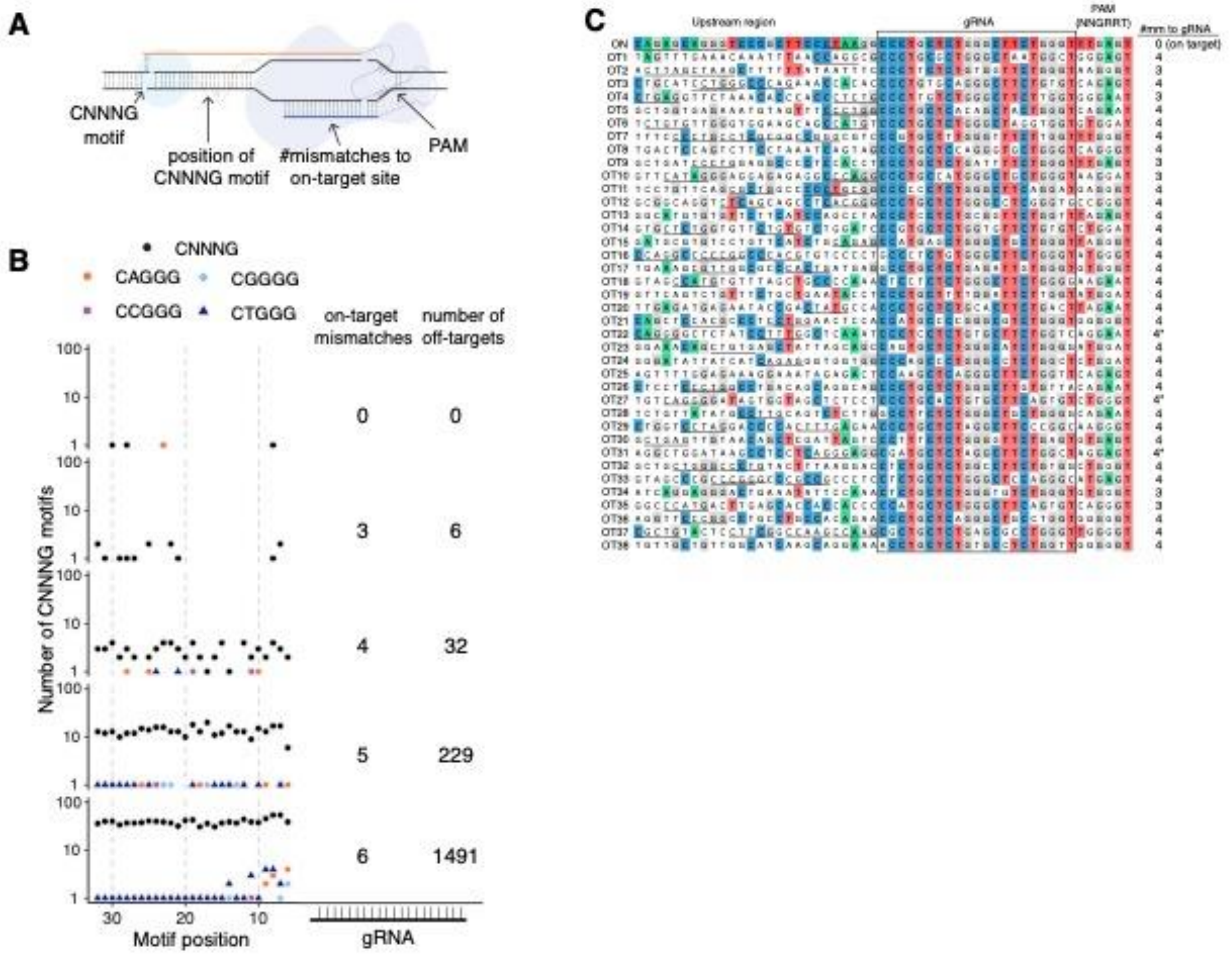


**Fig. S7. Identification of *AAVS1* gRNA off-targets in the human genome. A**, Schematic of methodology used to identify TevSaCas9 off-target sites. **B**, Plot of distribution of CNNNG motifs upstream of *AAVS1*-T1 off-target sites with the indicated mismatches to the gRNA portion. CNNNG that could support repair using the *AAVS1*-T1 rep-gRNA 3'-GG end are highlighted. C, Alignment of off-target sites to the on-target *AAVS1*-T1 sites. Identical nucleotides are colored, and CNNNG motifs in the upstream region are underlined.

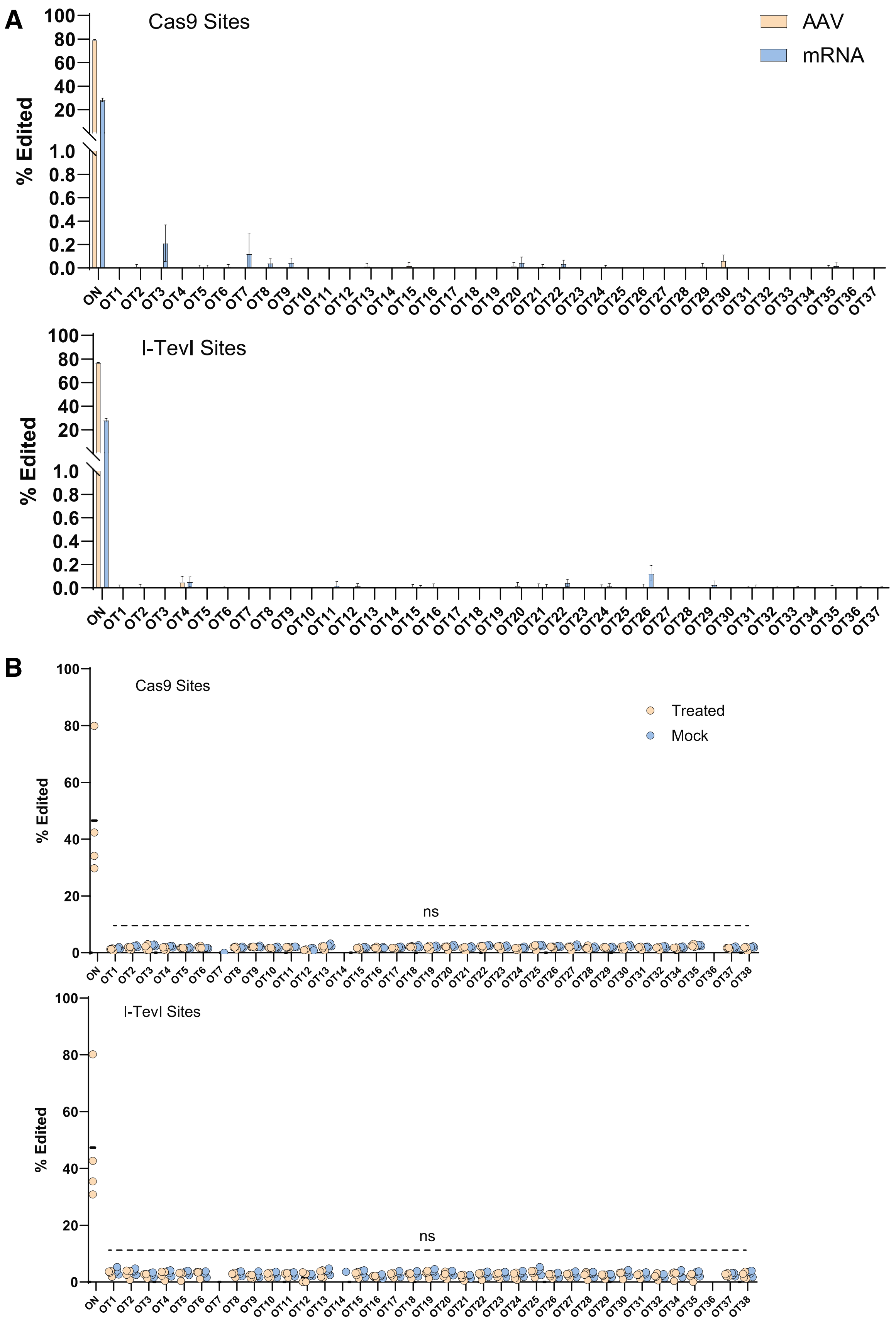
A
Cas9 Sites
AAV
mRNA
% Edited
I-TevI Sites
B
Treated
Mock
ns

**Fig. S8. A,** Percent editing at on- and off-target sites as determined by CRISPECTOR analysis for HEK293 treated with TevSaCas9/*AAVS1* rep-gRNA by AAV transduction or mRNA lipofection. The plots are separated into indels mapped to near the predicted SaCas9 cleavage site and Tev cleavage site for each target site. Bars are the mean of three replicates with whiskers representing 95% confidence interval from the CRISPECTOR calculated editing rate. **B**, Percent editing as determined by CRISPRaltRations analysis. Labeled as in panel A with points representing individual experiments. ns = not significant as calculated by paired t-tests of treated

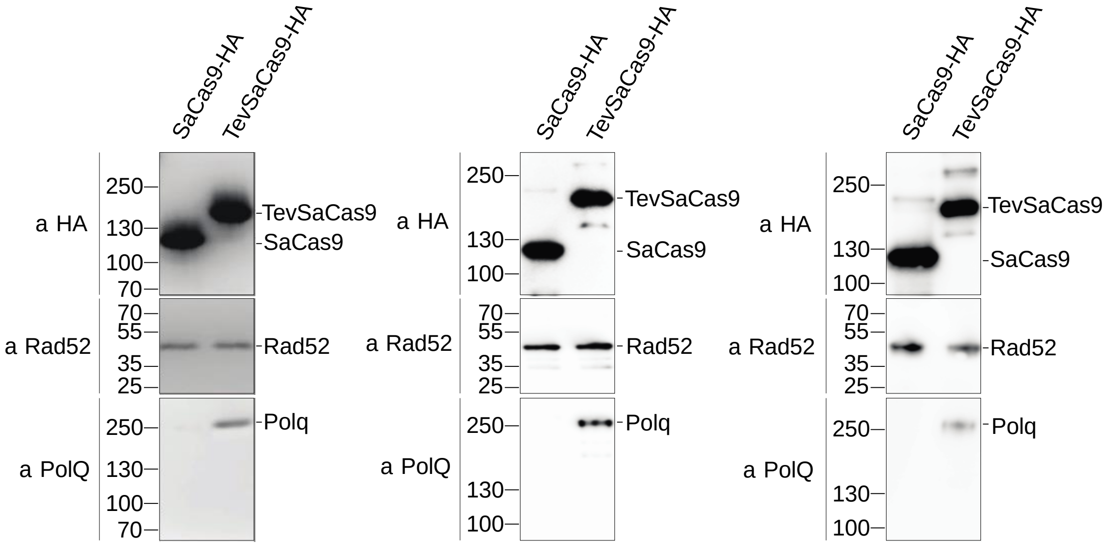


**Fig. S9. Interaction of Polq and Rad52 with TevSaCas9**. Shown are biological replicates (on left and right) of the co-immunoprecipitation experiment from HEK293 cells transfected with TevSsCas9/rep-gRNA or SaCas9/rep-gRNA shown in Fig. 2B (which is shown again here as the middle panel).

**Table. S1.**

Sequences of rep-gRNAs used in this study. All sequences are written as DNA in the 5’ to 3’ direction.

| Name | Target site (5’-3’) | Scaffold sequence (5’-3) | Repair template (5’-3’) | overhang | |
|---|---|---|---|---|---|
| | | | | 5` | 3’ |
| | *AAVS1*-related rep-gRNA | | | | |
| ***AAVS1*-T1-0** | CCCTGCTCTGGGCTTCTGGGT | GTTTTAGTACTCTGGAAACAGAATCTACTAAAACAAGGCAAAATGCCGTGTTTATCTCGTCAACTTGTTGGCGAGAT | AGACGAATAGAGAAGagatctGGAAAGCAATAAGA | 0 | 0 |
| ***AAVS1*-T1** | CCCTGCTCTGGGCTTCTGGGT | GTTTTAGTACTCTGGAAACAGAATCTACTAAAACAAGGCAAAATGCCGTGTTTATCTCGTCAACTTGTTGGCGAGAT | AGACGAATAGAGAAGagatctGGAAAGCAATAAGACC | 0 | 2 |
| ***AAVS1*-T1-+3+2** | CCCTGCTCTGGGCTTCTGGGT | GTTTTAGTACTCTGGAAACAGAATCTACTAAAACAAGGCAAAATGCCGTGTTTATCTCGTCAACTTGTTGGCGAGAT | ACCAGACGAATAGAGAAGagatctGGAAAGCAATAAGACC | 3 | 2 |
| ***AAVS1*-T1-+6+2** | CCCTGCTCTGGGCTTCTGGGT | GTTTTAGTACTCTGGAAACAGAATCTACTAAAACAAGGCAAAATGCCGTGTTTATCTCGTCAACTTGTTGGCGAGAT | CAAACCAGACGAATAGAGAAGagatctGGAAAGCAATAAGACC | 6 | 2 |
| ***AAVS1*-T1-+4** | CCCTGCTCTGGGCTTCTGGGT | GTTTTAGTACTCTGGAAACAGAATCTACTAAAACAAGGCAAAATGCCGTGTTTATCTCGTCAACTTGTTGGCGAGAT | AGACGAATAGAGAAGagatctGGAAAGCAATAAGACCTG | 0 | 4 |
| ***AAVS1*-T1-+6** | CCCTGCTCTGGGCTTCTGGGT | GTTTTAGTACTCTGGAAACAGAATCTACTAAAACAAGGCAAAATGCCGTGTTTATCTCGTCAACTTGTTGGCGAGAT | AGACGAATAGAGAAGagatctGGAAAGCAATAAGACCTGCT | 0 | 6 |
| ***AAVS1*-T1-+8** | CCCTGCTCTGGGCTTCTGGGT | GTTTTAGTACTCTGGAAACAGAATCTACTAAAACAAGGCAAAATGCCGTGTTTATCTCGTCAACTTGTTGGCGAGAT | AGACGAATAGAGAAGagatctGGAAAGCAATAAGACCTGCTCT | 0 | 8 |
| ***AAVS1*-T1-+10** | CCCTGCTCTGGGCTTCTGGGT | GTTTTAGTACTCTGGAAACAGAATCTACTAAAACAAGGCAAAATGCCGTGTTTATCTCGTCAACTTGTTGGCGAGAT | AGACGAATAGAGAAGagatctGGAAAGCAATAAGACCTGCTCTGG | 0 | 10 |
| ***AAVS1*-T1-+12** | CCCTGCTCTGGGCTTCTGGGT | GTTTTAGTACTCTGGAAACAGAATCTACTAAAACAAGGCAAAATGCCGTGTTTATCTCGTCAACTTGTTGGCGAGAT | AGACGAATAGAGAAGagatctGGAAAGCAATAAGACCTGCTCTGGGC | 0 | 12 |
| ***AAVS1*-T1-+14** | CCCTGCTCTGGGCTTCTGGGT | GTTTTAGTACTCTGGAAACAGAATCTACTAAAACAAGGCAAAATGCCGTGTTTATCTCGTCAACTTGTTGGCGAGAT | AGACGAATAGAGAAGagatctGGAAAGCAATAAGACCTGCTCTGGGCGG | 0 | 14 |
| ***AAVS1*-T1-+16** | CCCTGCTCTGGGCTTCTGGGT | GTTTTAGTACTCTGGAAACAGAATCTACTAAAACAAGGCAAAATGCCGTGTTTATCTCGTCAACTTGTTGGCGAGAT | AGACGAATAGAGAAGagatctGGAAAGCAATAAGACCTGCTCTGGGCGGAG | 0 | 16 |
| ***AAVS1*-T1-+18** | CCCTGCTCTGGGCTTCTGGGT | GTTTTAGTACTCTGGAAACAGAATCTACTAAAACAAGGCAAAATGCCGTGTTTATCTCGTCAACTTGTTGGCGAGAT | AGACGAATAGAGAAGagatctGGAAAGCAATAAGACCTGCTCTGGGCGGAGGA | 0 | 18 |

| | | | | | |
|---|---|---|---|---|---|
| ***AAVS1*-D13** | CCCTGCT CTGGGCT TCTGGGT | GTTTTAGTACTCTGGAAACAGAATCTACTAAAA CAAGGCAAAATGCCGTGTTTATCTCGTCAACT TGTTGGCGAGAT | CAGAAGCTTAGGGAAGCG GGACCC | 0 | 2 |
| | *CFTR*-related rep-gRNAs | | | | |
| **F508del** | ATATCATT GGTGTTT CCTATG | GTTTTAGTACTCTGGAAACAGAATCTA CTAAAACAAGGCAAAATGCCGTGTTT ATCTCGTCAACTTGTTGGCGAGAT | AGctcACgCCAaagATaATATT TTCTTTAATGGTGCCA | 0 | 2 |
| **G542X** | CTTtGAG AAGGTGG AATCACA | GTTTTAGTACTCTGGAAACAGAATCTA CTAAAACAAGGCAAAATGCCGTGTTT ATCTCGTCAACTTGTTGGCGAGAT | aATgCCgCCTTCTCcAAGAA CaATATTGTCTTTCtc | 0 | 2 |
| **W1282X** | TCCTtCA CTGTTGC AAAGTTA | GTTTTAGTACTCTGGAAACAGAATCTA CTAAAACAAGGCAAAATGCCGTGTTT ATCTCGTCAACTTGTTGGCGAGAT | CccTcCAgCAGTGgAGGAAA GCtTTTGGAGTGATACCac | 0 | 2 |
| | *SERPINA1*-related rep-gRNAs | | | | |
| ***SERPIN A1* E342K** | TGACCAT CGACaAG AAAGGGA | GTTTTAGTACTCTGGAAACAGAATCTA CTAAAACAAGGCAAAATGCCGTGTTT ATCTCGTCAACTTGTTGGCGAGAT | CcTTtTCaTCaATtGTCAGCA CAGCCTTATGCACGGCCT | 0 | 2 |

**Table S2.**

Oligonucleotides used in this study. All sequences are DNA and written in the 5' to 3' direction.

| Oligos ID | Sequence[5'-3'] | Modification | Note |
|---|---|---|---|
| Genomic PCR oligos | | | |
| OL0345 | GCAGCTTCCTTACACTTCCC | | gDNA *AAVS1* forward |
| OL0346 | CTAGGACTGAGGGTTTCAGTGC | | gDNA *AAVS1* reverse |
| OL1026 | TAGATCCTGCAGGGGTGTGA | | gDNA *CFTR* G542X forward |
| OL1027 | GACCTCCTCCCCCAACAATG | | gDNA *CFTR* G542X revere |
| OL1286 | GGTTCCTCCCCTGTGATTCC | | gDNA *SERPINA1* forward |
| OL1287 | GGCGACCAATGAACAACTGC | | gDNA *SERPINA1* reverse |
| OL0560 | ACACTCTTTCCCTACACGACGCTCTTCCGATCTCCTTTGTGAGAATGGTGCGTCC | | NGS *AAVS1* forward and RT-PCR |
| OL0561 | GACTGGAGTTCAGACGTGTGCTCTTCCGATCTGGCTCAGTCTGAAGAGCAGAGC | | NGS *AAVS1* reverse and RT-PCR |
| OL1543 | GATTACATTAGAAGGAAGATGTGC | | NGS *CFTR* G542X forward |
| OL1544 | ACAGCAAATGCTTGCTAGA | | NGS *CFTR* G542X reverse |
| OL1290 | TCGTCGGCAGCGTCAGATGTGTATAAGAGACAGGGGACCAGCTCAACCCTTCT | | NGS *SERPINA1* forward |
| OL1291 | GTCTCGTGGGCTCGGAGATGTGTATAAGAGACAGGAGGCGCTTGTCAGGAAGAT | | NGS *SERPINA1* reverse |
| RT-PCR oligos | | | |
| OL2504 | AGACGAATAGAGAAGag | 3'Phos | screen correct *AAVS1* repair and RT-PCR |
| OL2505 | ACCTCTTATTGCTTTCCag | 3'Phos | screen wrong *AAVS1* repair and RT-PCR |
| OL1081 | AGAAGGGCTGCCACAGCAGCATC | | hTev Forward primer RT-PCR |
| OL1082 | GATGATCTCCTCCTTCAG | | hTev Reverse primer RT-PCR |
| OL1083 | ACTACGAGACACGGGACGTG | | SaCas9 Forward primer RT-PCR |
| OL1084 | AGCAGGTTGTAGTCGAACAG | | Sacas9 Reverse primer RT-PCR |
| OL1085 | ACGGCCACAAGTTCAGCGTGTC | | eGFP Forward primer RT-PCR |

| | | | |
|---|---|---|---|
| OL1086 | TAGCGGCTGAAGCACTGCAC | | eGFP Reverse primer RT-PCR |
| OL1087 | CACCATTGGCAATGAGCGGTTC | | B-Actin Forward primer RT-PCR |
| OL1088 | AGGTCTTTGCGGATGTCCACGT | | B-Actin Reverse primer RT-PCR |
| rep-gRNA synthesis oligos | | | |
| OL2512 | aagcTAATACGACTCACTATAGGCCCTGCTCTGGGCTTCTGGGTGTTTTA GTACTCTGGAAACAGAATCTACTAAAACAAGGCAAAATGC | | T7-*AAVS1*-scaffold Klenow forward |
| OL2513 | GGTCTTATTGCTTTCCagatctCTTCTCTATTCGTCTTCTCGCCAACAAGTT GACGAGATAAACACGGCATTTTGCCTTGTTTTAGTAG | | *AAVS1* RT CA v1.0 Scaffold Klenow reverse |
| OL2514 | GGTCTTATTGCTTTCCagatctCTTCTCTATTCGTCTGGTTCTCGCCAACA AGTTGACGAGATAAACACGGCATTTTGCCTTGTTTTA | | *AAVS1* RT CA v2.0 Scaffold Klenow reverse |
| OL2515 | GGTCTTATTGCTTTCCagatctCTTCTCTATTCGTCTGGTTTCTCTCGCCAA CAAGTTGACGAGATAAACACGGCATTTTGCCTTGTTTT | | *AAVS1* RT CA v3.0 Scaffold Klenow reverse |
| OL2516 | GGTCTTATTGCTTTCCagatc | | *AAVS1* RT CA v1.0 Scaffold reverse |
| OL2517 | TCTTATTGCTTTCCagatc | | *AAVS1* RT CA reverse v5 |
| OL2518 | CAGGTCTTATTGCTTTCCagatc | | *AAVS1* RT CA reverse v6 |
| OL2519 | AGCAGGTCTTATTGCTTTCCagatc | | *AAVS1* RT CA reverse v7 |
| OL2520 | AGAGCAGGTCTTATTGCTTTCCagatc | | *AAVS1* RT CA reverse v8 |
| OL2521 | CCAGAGCAGGTCTTATTGCTTTCCagatc | | *AAVS1* RT CA reverse v9 |
| OL2522 | GCCCAGAGCAGGTCTTATTGCTTTCCagatc | | *AAVS1* RT CA reverse v10 |
| OL2523 | CCGCCCAGAGCAGGTCTTATTGCTTTCCagatc | | *AAVS1* RT CA reverse v11 |
| OL2524 | TTCCGCCCAGAGCAGGTCTTATTGCTTTCCagatc | | *AAVS1* RT CA reverse v12 |
| OL2421 | aagcTAATACGACTCACTATAGATATCATTGGTGTTTCCTATGGTTTTAGTA CTCTGGAAACAGAATCTACTAAAACAAGGCAAAATGCCGTG | | T7-*CFTR* F508del rep-gRNA forward |
| OL2422 | TGGCACCATTAAAGAAAATATtATcttTGGcGTgagCTTCTCGCCAACAAG TTGACGAGATAAACACGGCATTTTGCCTTGTTTTAG | | *CFTR* F508 del rep-gRNA reverse |

| | | | |
|---|---|---|---|
| OL2425 | aagcTAATACGACTCACTATAGCTTtGAGAAGGTGGAATCACAGTTTTAGTACTCTGGAAACAGAATCTACTAAAACAAGGCAAAATGCCGTGTTTATC | | T7-*CFTR* G542X rep-gRNA forward |
| OL2426 | gaGAAAGACAATATtGTTCTTgGAGAAGGcGGcATtTCTCGCCAACAAGTTGACGAGATAAACACGGCATTTTGCCTTGTTTTAGTAG | | *CFTR* G542X rep-gRNA reverse |
| OL2429 | aagcTAATACGACTCACTATAGTCCTtCACTGTTGCAAAGTTAGTTTTAGTACTCTGGAAACAGAATCTACTAAAACAAGGCAAAATGCCGTGTTTATC | | T7-*CFTR* W1282X rep-gRNA forward |
| OL2430 | gtGGTATCACTCCAAAaGCTTTCCTcCACTGcTGgAggGTCTCGCCAACAAGTTGACGAGATAAACACGGCATTTTGCCTTGTTTTAGTAG | | *CFTR* W1282X rep-gRNA reverse |
| OL2146 | TCGTCAACTTGTTGGCGAGAGgAAaAGtAGtTAaCAGTCGTGTCGGAATAC | | T7-*SERPINA1* E342K rep-gRNA forward |
| OL2147 | ACAGATCTAATGAAAATAAAGATCTTTTATTCGGCACGTATTCCGACACGACTGtTAaC | | *SERPINA1* E342K rep-gRNA reverse |
| Cloning oligos | | | |
| OL2244 | gctcgctttcttgctgtccaatttctattaaagCTCGCTGATCAGCCTCGACTG | | Generate backbone |
| OL2203 | TTCGTCCGAAGGCGGACTCATCAGTcgagcgaaAGCACGCGCTCTACCACTGAG | | Generate backbone |
| OL2580 | CTGATGAGTCCGCCTTCGGACGAACCAGAATAACGATAGCATGTACGAGGTCCCGGGTTC | | . Generate insert from [A]tRNA' to gRNA scaffold<br>. Generate insert [A]tRNA'-*AAVS1* rep gRNA -2xHBB |
| OL1907 | TCTCGCCAACAAGTTGACGAGATAAACAC | | Generate insert from [A]tRNA' to gRNA scaffold |
| OL2581 | GTGTTTATCTCGTCAACTTGTTGGCGAGAAGACGAATAGAGAAGagatctGGAAAG | | Generate *AAVS1* RT CA Klenow |
| OL2582 | tggacagcaagaaagcgagcTGGTCTTATTGCTTTCCagatctCTTCTCTATTC | | . Generate *AAVS1* RTCA Klenow<br>. Generate insert [A]tRNA'-*AAVS1* rep_gRNA CA-2xHBB |
| OL0198 | CACCCCCTGCTCTGGGCTTCTGGGT | | Clone *AAVS1* gRNA sense |
| OL0199 | AAACACCCAGAAGCCCAGAGCAGGG | | Clone *AAVS1* gRNA antisense |
| OL2214 | CTGATGAGTCCGCCTTCGGACGAAgaGAAAGACAATTAGCATGTACGAGGTCCCGGGTTC | | Generate tRNA' specific for *CFTR* G542X rep gRNA |

| | | | |
|---|---|---|---|
| OL1907 | TCTCGCCAACAAGTTGACGAGATAAACAC | | Generate tRNA' specific for *CFTR* G542X rep gRNA |
| OL2439 | GTGTTTATCTCGTCAACTTGTTGGCGAGAaATgCCgCCTTCTCcAAGAACaATATTGTC | | Generate RT0052 CAv2.0 with overlap for *CFTR* G542X gRNA and 2xHBB |
| OL2445 | gaaattggacagcaagaaagcgagcTgaGAAAGACAATATtGTTCTTgGAGAAG | | Generate RT0052 CAv2.0 with overlap for *CFTR* G542X gRNA and 2xHBB |
| OL2411 | CTGATGAGTCCGCCTTCGGACGAAggccgtgcataaTAGCATGTACGAGGTCCCGGGTTC | | . Generate insert from [A]tRNA' to gRNA scaffold<br>. Generate insert [A]tRNA'-Serpina1 E342K rep gRNA CAv2.0-2xHBB |
| OL2148 | TCTCGTCAACTTGTTGGCGAGACcTTtTCaTCaATtGTCAGCACAGCCTTATG | | Generate Serpina1 E342K rep gRNACAv2.0 Klenow |
| OL2412 | tggacagcaagaaagcgagcAGGCCGTGCATAAGGCTGTGCTGACaATtG | | . Generate Serpina1 E342K rep gRNACAv2.0 Klenow<br>. Generate insert [A]tRNA'-Serpina1 E342K rep gRNA CAv2.0-2xHBB |
| OL2548 | GTTGGACCGGTGCCACCATGGCCGACGCCACCTTCGGCGACACCTG | | AgeI-Kozak-I-Tev[del1-92] forward |
| rep-gRNA specificity oligos | | | |
| OL2782 | aagcTAATACGACTCACTATAGGCACTGCTCTGGGCTTCTGGGTGTTTTAGTACTCTGGAAACAGAATCTACTAAAACAAGGCAAAATGC | | T7-*AAVS1* gRNA forward (KD) gv01 |
| OL2783 | aagcTAATACGACTCACTATAGGCCCTACTCTGGGCTTCTGGGTGTTTTAGTACTCTGGAAACAGAATCTACTAAAACAAGGCAAAATGC | | T7-*AAVS1* gRNA forward (KD) gv02 |
| OL2784 | aagcTAATACGACTCACTATAGGCCCTGCTATGGGCTTCTGGGTGTTTTAGTACTCTGGAAACAGAATCTACTAAAACAAGGCAAAATGC | | T7-*AAVS1* gRNA forward (KD) gv03 |
| OL2785 | aagcTAATACGACTCACTATAGGCCCTGCTCTGAGCTTCTGGGTGTTTTAGTACTCTGGAAACAGAATCTACTAAAACAAGGCAAAATGC | | T7-*AAVS1* gRNA forward (KD) gv04 |

| | | | |
|---|---|---|---|
| OL2786 | aagcTAATACGACTCACTATAGGCCCTGCTCTGGGCATCTGGGTGTTTT AGTACTCTGGAAACAGAATCTACTAAAACAAGGCAAAATGC | | T7-*AAVS1* gRNA forward (KD) gv05 |
| OL2787 | aagcTAATACGACTCACTATAGGCCCTGCTCTGGGCTTCAGGGTGTTTT AGTACTCTGGAAACAGAATCTACTAAAACAAGGCAAAATGC | | T7-*AAVS1* gRNA forward (KD) gv06 |
| OL2788 | aagcTAATACGACTCACTATAGGCCCTGCTCTGGGCTTCTGGATGTTTTA GTACTCTGGAAACAGAATCTACTAAAACAAGGCAAAATGC | | T7-*AAVS1* gRNA forward (KD) gv07 |
| OL2789 | aagcTAATACGACTCACTATAGGCACTACTCTGGGCTTCTGGGTGTTTTA GTACTCTGGAAACAGAATCTACTAAAACAAGGCAAAATGC | | T7-*AAVS1* gRNA forward (KD) gv08 |
| OL2790 | aagcTAATACGACTCACTATAGGCCCTGCTCTGAGCATCTGGGTGTTTTA GTACTCTGGAAACAGAATCTACTAAAACAAGGCAAAATGC | | T7-*AAVS1* gRNA forward (KD) gv09 |
| OL2791 | aagcTAATACGACTCACTATAGGCCCTGCTCTGGGCTTCAGGATGTTTTA GTACTCTGGAAACAGAATCTACTAAAACAAGGCAAAATGC | | T7-*AAVS1* gRNA forward (KD) gv10 |
| OL2792 | aagcTAATACGACTCACTATAGGCCCTACTCTGAGCTTCAGGGTGTTTTA GTACTCTGGAAACAGAATCTACTAAAACAAGGCAAAATGC | | T7-*AAVS1* gRNA forward (KD) gv11 |
| OL2793 | GTTCTTATTGCTTTCCagatctCTTCTCTATTCGTCTATCTCGCCAACAAGT TGACGAGATAAACACGGCATTTTGCCTTGTTTTAGTAG | | *AAVS1* RT-scaffold reverse KD v01 |
| OL2794 | GCTCTTATTGCTTTCCagatctCTTCTCTATTCGTCTATCTCGCCAACAAGT TGACGAGATAAACACGGCATTTTGCCTTGTTTTAGTAG | | *AAVS1* RT-scaffold reverse KD v02 |
| OL2795 | GATCTTATTGCTTTCCagatctCTTCTCTATTCGTCTATCTCGCCAACAAGT TGACGAGATAAACACGGCATTTTGCCTTGTTTTAGTAG | | *AAVS1* RT-scaffold reverse KD v03 |
| OL2796 | CCTCTTATTGCTTTCCagatctCTTCTCTATTCGTCTATCTCGCCAACAAGT TGACGAGATAAACACGGCATTTTGCCTTGTTTTAGTAG | | *AAVS1* RT-scaffold reverse KD v04 |
| OL2797 | CGTCTTATTGCTTTCCagatctCTTCTCTATTCGTCTATCTCGCCAACAAGT TGACGAGATAAACACGGCATTTTGCCTTGTTTTAGTAG | | *AAVS1* RT-scaffold reverse KD v05 |
| OL2798 | CTTCTTATTGCTTTCCagatctCTTCTCTATTCGTCTATCTCGCCAACAAGT TGACGAGATAAACACGGCATTTTGCCTTGTTTTAGTAG | | *AAVS1* RT-scaffold reverse KD v06 |

| | | | |
|---|---|---|---|
| OL2799 | CATCTTATTGCTTTCCagatctCTTCTCTATTCGTCTATCTCGCCAACAAGTTGACGAGATAAACACGGCATTTTGCCTTGTTTTAGTAG | | *AAVS1* RT-scaffold reverse KD v07 |
| OL2800 | TTTCTTATTGCTTTCCagatctCTTCTCTATTCGTCTATCTCGCCAACAAGTTGACGAGATAAACACGGCATTTTGCCTTGTTTTAGTAG | | *AAVS1* RT-scaffold reverse KD v08 |
| OL2801 | TGTCTTATTGCTTTCCagatctCTTCTCTATTCGTCTATCTCGCCAACAAGTTGACGAGATAAACACGGCATTTTGCCTTGTTTTAGTAG | | *AAVS1* RT-scaffold reverse KD v09 |
| OL2802 | TCTCTTATTGCTTTCCagatctCTTCTCTATTCGTCTATCTCGCCAACAAGTTGACGAGATAAACACGGCATTTTGCCTTGTTTTAGTAG | | *AAVS1* RT-scaffold reverse KD v10 |
| OL2803 | TATCTTATTGCTTTCCagatctCTTCTCTATTCGTCTATCTCGCCAACAAGTTGACGAGATAAACACGGCATTTTGCCTTGTTTTAGTAG | | *AAVS1* RT-scaffold reverse KD v11 |
| OL2804 | AATCTTATTGCTTTCCagatctCTTCTCTATTCGTCTATCTCGCCAACAAGTTGACGAGATAAACACGGCATTTTGCCTTGTTTTAGTAG | | *AAVS1* RT-scaffold reverse KD v12 |
| OL2805 | AGTCTTATTGCTTTCCagatctCTTCTCTATTCGTCTATCTCGCCAACAAGTTGACGAGATAAACACGGCATTTTGCCTTGTTTTAGTAG | | *AAVS1* RT-scaffold reverse KD v13 |
| OL2806 | ACTCTTATTGCTTTCCagatctCTTCTCTATTCGTCTATCTCGCCAACAAGTTGACGAGATAAACACGGCATTTTGCCTTGTTTTAGTAG | | *AAVS1* RT-scaffold reverse KD v14 |
| OL2807 | ATTCTTATTGCTTTCCagatctCTTCTCTATTCGTCTATCTCGCCAACAAGTTGACGAGATAAACACGGCATTTTGCCTTGTTTTAGTAG | | *AAVS1* RT-scaffold reverse KD v15 |

**Table S3.**
List of mRNAs used in this study.

| mRNA ID | 5'UTR | Key RNA region | 3' UTR | Target gene | Target allele |
|---|---|---|---|---|---|
| MR0011 | Human hemoglobin subunit beta | I-TevI[VKN]-SaCas9[WT]-2xNuc NLS | 2x human hemoglobin subunit beta | N/A | N/A |
| MR0012 | Human hemoglobin subunit beta | SaCas9[WT]-2xNuc NLS | 2x human hemoglobin subunit beta | N/A | N/A |
| MR0013 | Hybrid intron | I-TevI[VKN]-SaCas9[WT]-2xNuc NLS-T2A-eGFP-MALAT-tRNA[HHRz]-*AAVS1* rep_gRNA | 2x human hemoglobin subunit beta | *AAVS1* | *AAVS1* wildtype |
| MR0020 | Tobacco Etch Virus 5' Leader sequence | I-TevI[VKN]-SaCas9[WT]-2xNuc NLS-T2A-eGFP-MALAT-tRNA[HHRz]-*AAVS1* rep_gRNA | 2x human hemoglobin subunit beta | *AAVS1* | *AAVS1* wildtype |
| MR0059 | Tobacco Etch Virus 5' Leader sequence | I-TevI[VKN]-SaCas9[WT]-2xNuc NLS-T2A-eGFP-MALAT-tRNA[HHRz]-Serpina1 E342K rep_gRNA | 2x human hemoglobin subunit beta | Serpina1 | E342K |
| MR0086 | Tobacco Etch Virus 5' Leader sequence | I-TevI[VKN]-SaCas9[WT]-2xNuc NLS-T2A-eGFP-MALAT-tRNA[HHRz]-*CFTR* G542X rep_gRNA | 2x human hemoglobin subunit beta | *CFTR* | G542X |

**Table S4.**
List of AAV viral vectors used in this study.

| AAV ID | Promoter | Key RNA region | Serotype | Target gene | Target allele |
|---|---|---|---|---|---|
| AAV0005 | mini CMV | I-TevI[VKN]-SaCas9[WT]-HH-*AAVS1* gRNA-HDV-SV40[A] | 2 | *AAVS1* | *AAVS1* wildtype |
| AAV0024 | mini CMV | I-TevI[VKN]-SaCas9[WT]-SV40 NLS-MALAT-tRNA[HHRz]-*AAVS1* rep_gRNA-Synt[A] | 9 | *AAVS1* | *AAVS1* wildtype |
| AAV0026 | mini CMV | I-TevI[VKN]-SaCas9[WT]-SV40 NLS-MALAT-tRNA[HHRz]-*AAVS1* rep_gRNA-Synt[A] | 2 | *AAVS1* | *AAVS1* wildtype |

**Table S5.**
*AAVS1*-T1 gRNA off-targets.

| crRNA | DNA | Xosome | Position | Direction | Mismatches | Bulge Size |
|---|---|---|---|---|---|---|
| CCCTGCTCTGGGCTTCTGGGTNNGRRT | CCCTGCgCTGGGCTaaTGGcTGGGAGT | chr5 | 1133265 | + | 4 | 0 |
| CCCTGCTCTGGGCTTCTGGGTNNGRRT | CCCTtCTCTGtGgTTCTGGGTAAGGGT | chr5 | 58205673 | - | 3 | 0 |
| CCCTGCTCTGGGCTTCTGGGTNNGRRT | CCCTGtgCaGGGCTTCTGtGTCAGAGT | chr5 | 170525900 | + | 4 | 0 |
| CCCTGCTCTGGGCTTCTGGGTNNGRRT | CCCTtgTCTGGGCTTCTtGGTGGGAAT | chr5 | 172545533 | + | 3 | 0 |
| CCCTGCTCTGGGCTTCTGGGTNNGRRT | CCCTGCTCacaGCTaCTGGGTCAGAAT | chr20 | 23607012 | - | 4 | 0 |
| CCCTGCTCTGGGCTTCTGGGTNNGRRT | CCCTGCTCTGGGCTaggtGGTGTGGAT | chr20 | 25507534 | - | 4 | 0 |
| CCCTGCTCTGGGCTTCTGGGTNNGRRT | CCgTGCTtTGGGtTTCTtGGTTTGGGT | chr20 | 63956973 | - | 4 | 0 |
| CCCTGCTCTGGGCTTCTGGGTNNGRRT | CCCTGCTCcaGGgTgCTGGGTCAGGGT | chr1 | 17733177 | + | 4 | 0 |
| CCCTGCTCTGGGCTTCTGGGTNNGRRT | CCCTGCTCTGattTTCTGGGTTTGAGT | chr1 | 22617684 | + | 3 | 0 |
| CCCTGCTCTGGGCTTCTGGGTNNGRRT | CCCTGCcaTGGGCTgCTGGGTAAGGAT | chr1 | 30173728 | + | 3 | 0 |
| CCCTGCTCTGGGCTTCTGGGTNNGRRT | CCCccCTCTGGGCTTCaGGaTGAGGGT | chr22 | 44671178 | - | 4 | 0 |
| CCCTGCTCTGGGCTTCTGGGTNNGRRT | CCCTGCTCTGGGCcTCgGGtgCCGGGT | chr7 | 1478934 | - | 4 | 0 |
| CCCTGCTCTGGGCTTCTGGGTNNGRRT | CCCTcCTCTGcGgTTCTGGtTTAGAGT | chr7 | 154318967 | + | 4 | 0 |
| CCCTGCTCTGGGCTTCTGGGTNNGRRT | CCgTGCTCTGGtgTTCTGtGTCTGGAT | chr2 | 3478135 | + | 4 | 0 |
| CCCTGCTCTGGGCTTCTGGGTNNGRRT | CCaTGagCTGGGCTgCTGGGTTAGGGT | chr2 | 104816468 | - | 4 | 0 |
| CCCTGCTCTGGGCTTCTGGGTNNGRRT | gCCctCTgTGGGCTTCTGGGTATGGGT | chr12 | 54146864 | - | 4 | 0 |

| | | | | | | |
|---|---|---|---|---|---|---|
| CCCTGCTCTGGGCTTCTGGGTNNGRRT | gCCTGCTCTGaGaTTgTGGGTGTGGGT | chr17 | 45900070 | + | 4 | 0 |
| CCCTGCTCTGGGCTTCTGGGTNNGRRT | CtCctCTCTGGGCTTCTGGGgAAGAAT | chr17 | 57880611 | - | 4 | 0 |
| CCCTGCTCTGGGCTTCTGGGTNNGRRT | CCCTGCTtTtGGaTTCTtGGTATGGAT | chr17 | 66221697 | + | 4 | 0 |
| CCCTGCTCTGGGCTTCTGGGTNNGRRT | CCCTGCTCTGcaCTTCTGacTTAGAAT | chr17 | 66729035 | - | 4 | 0 |
| CCCTGCTCTGGGCTTCTGGGTNNGRRT | CCaTGCcCcGGGCgTCTGGGTGGGGGT | chr17 | 75742569 | - | 4 | 0 |
| CCCTGCTCTGGGCTTCTGGGTNNGRRT | CCCTcCTCTGtGCTTCTGGtcAGGAAT | chr16 | 20541514 | + | 4 | 0 |
| CCCTGCTCTGGGCTTCTGGGTNNGRRT | CagTGCTCTGGGCaTCTGGGgATGGAT | chr16 | 24497213 | - | 4 | 0 |
| CCCTGCTCTGGGCTTCTGGGTNNGRRT | CCCaGCcCTGGGCcTCTGGcTCTGGAT | chr16 | 55003484 | + | 4 | 0 |
| CCCTGCTCTGGGCTTCTGGGTNNGRRT | CCaaGCTCaGGGCTTCTGGtTCAGAGT | chr16 | 65201928 | + | 4 | 0 |
| CCCTGCTCTGGGCTTCTGGGTNNGRRT | CCCTGCTCTGGGCTTgTGttaCAGAAT | chr9 | 35391588 | - | 4 | 0 |
| CCCTGCTCTGGGCTTCTGGGTNNGRRT | CCCTGCaCTGtGCTTCaGtGTCTGGGT | chrX | 44391139 | - | 4 | 0 |
| CCCTGCTCTGGGCTTCTGGGTNNGRRT | gCCTtCTCTGGGCTgCTGGGgCAGAAT | chr14 | 88308410 | - | 4 | 0 |
| CCCTGCTCTGGGCTTCTGGGTNNGRRT | CCCTGCTCTaGGCTTCccGGcAAGGGT | chr6 | 75930672 | - | 4 | 0 |
| CCCTGCTCTGGGCTTCTGGGTNNGRRT | CCtTtCTCTGGGgTTCTGaGTGTGAGT | chr6 | 90764834 | + | 4 | 0 |
| CCCTGCTCTGGGCTTCTGGGTNNGRRT | CgaTGCTCTaGGCTTCTGGcTAGGAGT | chr10 | 43959582 | - | 4 | 0 |
| CCCTGCTCTGGGCTTCTGGGTNNGRRT | CtCTGCTCTGGcCTTCTGtGgCTGGGT | chr13 | 25863207 | + | 4 | 0 |
| CCCTGCTCTGGGCTTCTGGGTNNGRRT | CtCTGCTCTGGGCTcCaGGGcATGAGT | chr13 | 33350183 | - | 4 | 0 |
| CCCTGCTCTGGGCTTCTGGGTNNGRRT | CtCTGCTCTGGGtgTCTGGGTGTGGGT | chr13 | 103148432 | + | 3 | 0 |

| | | | | | | |
|---|---|---|---|---|---|---|
| CCCTGCTCTGGGCTTCTGGGTNNGRRT | CCaTGCTCTGGGCTTCaGtGTCAGGGT | chr19 | 11175767 | + | 3 | 0 |
| CCCTGCTCTGGGCTTCTGGGTNNGRRT | CCCTGCTCaGGGCTgCctGGTGGGGGT | chr19 | 19547992 | + | 4 | 0 |
| CCCTGCTCTGGGCTTCTGGGTNNGRRT | CgCTGCTCTGaGCgcCTGGGTTGGGGT | chr19 | 46630813 | + | 4 | 0 |
| CCCTGCTCTGGGCTTCTGGGTNNGRRT | CCCTGCTCTGGGCTTCTGGGTTTGAGT | chr19 | 55115040 | - | 0 | 0 |
| CCCTGCTCTGGGCTTCTGGGTNNGRRT | aCCTGCTCTGtGCcTCTGGtTGGGGGT | chr3 | 10920852 | + | 4 | 0 |

### References


1. H. C. Valley, K. M. Bukis, A. Bell, Y. Cheng, E. Wong, N. J. Jordan, N. E. Allaire, A. Sivachenko, F. Liang, H. Bihler, others, Isogenic cell models of cystic fibrosis-causing variants in natively expressing pulmonary epithelial cells. *Journal of Cystic Fibrosis* **18**, 476–483 (2019).
2. F. A. Ran, L. Cong, W. X. Yan, D. A. Scott, J. S. Gootenberg, A. J. Kriz, B. Zetsche, O. Shalem, X. Wu, K. S. Makarova, others, In vivo genome editing using Staphylococcus aureus Cas9. *Nature* **520**, 186–191 (2015).
3. N. Huynh, J. Zeng, W. Liu, K. King-Jones, A Drosophila CRISPR/Cas9 toolkit for conditionally manipulating gene expression in the prothoracic gland as a test case for polytene tissues. *G3: Genes, Genomes, Genetics* **8**, 3593–3605 (2018).
4. D. Kostyushev, A. Kostyusheva, S. Brezgin, D. Zarifyan, A. Utkina, I. Goptar, V. Chulanov, Suppressing the NHEJ pathway by DNA-PKcs inhibitor NU7026 prevents degradation of HBV cccDNA cleaved by CRISPR/Cas9. *Sci Rep* **9**, 1847 (2019).
5. W.-W. Zhang, P. Lypaczewski, G. Matlashewski, Optimized CRISPR-Cas9 genome editing for Leishmania and its use to target a multigene family, induce chromosomal translocation, and study DNA break repair mechanisms. *mSphere* **2**, 10–1128 (2017).
6. T. Killian, S. Dickopf, A. K. Haas, C. Kirstenpfad, K. Mayer, U. Brinkmann, Disruption of diphthamide synthesis genes and resulting toxin resistance as a robust technology for quantifying and optimizing CRISPR/Cas9-mediated gene editing. *Sci Rep* **7**, 15480 (2017).
7. F. Huang, N. Goyal, K. Sullivan, K. Hanamshet, M. Patel, O. M. Mazina, C. X. Wang, W. F. An, J. Spoonamore, S. Metkar, others, Targeting BRCA1-and BRCA2-deficient cells with RAD52 small molecule inhibitors. *Nucleic Acids Res* **44**, 4189–4199 (2016).
8. G. Rodriguez-Berriguete, M. Ranzani, R. Prevo, R. Puliyadi, N. Machado, H. R. Bolland, V. Millar, D. Ebner, M. Boursier, A. Cerutti, others, Small-molecule Polθ inhibitors provide safe and effective tumor radiosensitization in preclinical models. *Clinical Cancer Research* **29**, 1631–1642 (2023).
9. N. Huynh, Q. Ou, P. Cox, R. Lill, K. King-Jones, Glycogen branching enzyme controls cellular iron homeostasis via Iron Regulatory Protein 1 and mitoNEET. *Nat Commun* **10**, 5463 (2019).
10. J. P. Connelly, S. M. Pruett-Miller, CRIS. py: a versatile and high-throughput analysis program for CRISPR-based genome editing. *Sci Rep* **9**, 4194 (2019).
11. K. Clement, H. Rees, M. C. Canver, J. M. Gehrke, R. Farouni, J. Y. Hsu, M. A. Cole, D. R. Liu, J. K. Joung, D. E. Bauer, others, CRISPResso2 provides accurate and rapid genome editing sequence analysis. *Nat Biotechnol* **37**, 224–226 (2019).
12. T. Magoč, S. L. Salzberg, FLASH: fast length adjustment of short reads to improve genome assemblies. *Bioinformatics* **27**, 2957–2963 (2011).
13. S. Bae, J. Park, J.-S. Kim, Cas-OFFinder: a fast and versatile algorithm that searches for potential off-target sites of Cas9 RNA-guided endonucleases. *Bioinformatics* **30**, 1473–1475 (2014).
14. J. M. Wolfs, T. A. Hamilton, J. T. Lant, M. Laforet, J. Zhang, L. M. Salemi, G. B. Gloor, C. Schild-Poulter, D. R. Edgell, Biasing genome-editing events toward precise length

deletions with an RNA-guided TevCas9 dual nuclease. *Proc Natl Acad Sci U S A* **113** (2016).

15. I. Amit, O. Iancu, A. Levy-Jurgenson, G. Kurgan, M. S. McNeill, G. R. Rettig, D. Allen, D. Breier, N. Ben Haim, Y. Wang, L. Anavy, A. Hendel, Z. Yakhini, CRISPECTOR provides accurate estimation of genome editing translocation and off-target activity from comparative NGS data. *Nature Communications 2021 12:1* **12**, 1–11 (2021).
16. G. Kurgan, R. Turk, H. Li, N. Roberts, G. R. Rettig, A. M. Jacobi, L. Tso, M. Sturgeon, M. Mertens, R. Noten, K. Florus, M. A. Behlke, Y. Wang, M. S. McNeill, CRISPAltRations: a validated cloud-based approach for interrogation of double-strand break repair mediated by CRISPR genome editing. *Mol Ther Methods Clin Dev* **21**, 478–491 (2021).
17. E. K. Brinkman, T. Chen, M. Amendola, B. Van Steensel, Easy quantitative assessment of genome editing by sequence trace decomposition. *Nucleic Acids Res* **42** (2014).
